# *Cerebellin-2* neurons in the parafascicular thalamic nucleus regulate self-grooming behavior

**DOI:** 10.1101/2025.03.22.644777

**Authors:** Li Zhang, Huimeng Song, Yaning Li, Huaxin Zhang, Shibo Diao, Dandan Geng, Zijie Zhao, Bo Yang, Rong Zheng, Zekun Li, Lei Wang, Tianyun Zhang, Yingyue Jiang, Zining Liu, Lexuan Zhang, Yihe Liu, Zhiyong Xie, Dapeng Li, Fan Zhang, Peng Cao

**Affiliations:** National Institute of Biological Sciences, Beijing, China. Tsinghua Institute of Multidisciplinary Biomedical Research, Tsinghua University, Beijing, China; School of Basic Medical Sciences, Laboratory for Clinical Medicine, Capital Medical University, Beijing, China; The Key Laboratory of Neural and Vascular Biology, Ministry of Education; The Key Laboratory of Vascular Biology of Hebei Province; Department of Biochemistry and Molecular Biology, Hebei Medical University, Shijiazhuang, China; Department of Psychological Medicine, Zhongshan Hospital, Institute for Translational Brain Research, State Key Laboratory of Medical Neurobiology, MOE Frontiers Center for Brain Science, MOE Innovative Center for New Drug Development of Immune Inflammatory Diseases, Fudan University, Shanghai, China

## Abstract

Self-grooming represents a stereotyped repetitive behavior which is commonly observed in various neuropsychiatric disorders. Deep brain stimulation (DBS) targeting the centromedian / parafascicular complex (CM/PF) has been shown to alleviate obsessive-compulsive disorder/behavior (OCD/OCB) symptoms in human. However, little is known about the neural circuits of the PF that are involved in the regulation of self-grooming behavior in rodents. Here, we report that *cerebellin-2* positive (*Cbln2*^+^) neurons in PF bidirectionally encode self-grooming. Chemogenetic activation of *Cbln2*^+^ PF neurons significantly reduces both physiological and pathological self-grooming, and chemogenetic inhibition of *Cbln2*^+^ PF neurons increases physiological self-grooming. Moreover, activating synaptic excitatory inputs to *Cbln2*^+^ PF neurons inhibits self-grooming, while synaptic inhibitory inputs enhance it. *Cbln2*^+^ PF neurons independently transmit neural signals to the dorsal and ventral striatum. Activation of PF-dorsal striatum and PF-ventral striatum pathways both inhibit self-grooming behavior. Most excitingly, we alleviate excessive grooming in OCD mice by the activation of PF neurons using non-invasive ultrasound stimulation. Together, these data reveal that *Cbln2*^+^ PF neurons are integral components of a brain network involved in self-grooming behavior and highly promising targets in the non-invasive therapy of OCD.

## INTRODUCTION

Self-grooming is an evolutionarily conserved behavior observed in arthropods (Lefebvre, 1981; Richard and Dawkins, 1976), birds (Lefebvre, 1982; Lefebvre and Joly, 1982; van RHIJN, 1977), and mammals (Berridge, 1990; Colonnese et al., 1996). In rodents, self-grooming has a complex sequenced structure that consists of repeated stereotyped movements, which follows an innate cephalo-caudal progression, preceding by a postural cephalo-caudal transition from paw - head grooming to body grooming (Kalueff et al., 2007; Kalueff et al., 2016). Humans also exhibit repetitive patterned behaviors including excoriation (skin-picking) (Ekore and Ekore, 2021; Lochner et al., 2017) and trichotillomania (hair-pulling) (Grant and Chamberlain, 2016; Hoffman et al., 2021; Stein et al., 2006), which are in analogy with the stereotyped repetitive self-grooming behavior in rodents and share several common traits and brain circuits with rodents (Grant and Chamberlain, 2016; Graybiel and Rauch, 2000; Robbins et al., 2019; Robertson, 2000; Stein et al., 2008). Repetitive patterned behaviors in human frequently occur in several neural neuropsychiatric diseases including autism spectrum disorder (ASD) (Sharma et al., 2018), Tourette syndrome (Robertson, 2000), and obsessive-compulsive disorder (OCD) (Goodman et al., 2021), making the grooming behavior in rodents ideal to study the neuropsychiatric disorders (Liu et al., 2020; Peça et al., 2011; Welch et al., 2007).

Excessive self-grooming is frequently observed in genetic animal models of neuropsychiatric disorders, including obsessive compulsive disorder/behavior (OCD/OCB) (Piantadosi et al., 2024; Xue et al., 2022), Tourette’ syndrome (TS) (Liu et al., 2020), autism spectrum disorder (Peñagarikano et al., 2011; Silverman et al., 2010), and Rett syndrome (Chao et al., 2010; Gold et al., 2024). Thus, understanding the neural circuitry that regulates self-grooming could provide a new framework for the research of the development neuropsychiatric disorders.

The parafascicular thalamic (PF) nucleus in rodents, known as the counterpart of centromedian/ parafascicular complex (CM/PF) in primates, belongs to the intralaminar system of the thalamus (Langlois et al., 2010; Smith et al., 2004). The PF nucleus play a role in limbic and associative network functions (Fallon et al., 2023; Lacey et al., 2007), and send glutamatergic projections to the striatum, which is involved in the completion of sequential syntactic self-grooming chains (Cromwell and Berridge, 1996; Zhang et al., 2022). Deep brain stimulation (DBS) targeting the CM/PF has been shown to alleviate obsessive-compulsive disorder/behavior (OCD/OCB) symptoms in human (Servello et al., 2020). However, the neural circuit mechanism by which PF regulates repeated stereotyped behaviors, such as self-grooming, remains unclear. More importantly, whether non-invasive treatment, such as ultrasound stimulation of PF can modulate obsessive-compulsive disorder/behavior (OCD/OCB) symptoms is worth exploring (Darmani et al., 2022).

In the present study, we combined genetically encoded circuit analysis tools to identify a subset of PF neurons critical for both biological and pathological self-grooming in mice. *Cbln2*^+^ PF neurons bidirectionally encode self-grooming. Activating synaptic inhibitory inputs to *Cbln2*^+^ PF neurons enhances self-grooming, while synaptic excitatory inputs inhibit it. Moreover, *Cbln2*^+^ PF neurons regulate self-grooming behavior through both dorsal and ventral striatum. Non-invasive ultrasound stimulation efficiently activates PF neurons while reducing time spent for spontaneous grooming in both WT mice and OCD mice. Our data reveal that *Cbln2*^+^ PF neurons are integral components of a brain network involved in self-grooming behavior and ultrasound stimulation targeting the PF holds clinical therapeutic significance in OCD.

## RESULTS

### *Cbln2*^+^ PF neurons are required for the regulation of self-grooming

At the beginning of this study, we measured self-grooming by using a magnetic induction method (Inagaki et al., 2002; Xie et al., 2022) in parallel with video recording, this was modified from the methods described in previous work (Xie et al., 2022) (Fig. 1A). In previous study, the magnet was implanted in the forelimb to track the movements of forelimb during orofacial grooming stage, but it did not effectively indicate other types of grooming. Here, we implanted the magnet on the surface of the skull to track the movements of head during all types of self-grooming (Fig. 1A). The voltage threshold was calculated as before (mean ± 5 × SD) and used to identify the voltage pulses corresponding to the grooming behaviors. The magnetic inducing voltage signals efficiently indicated the grooming behaviors in each stage, including paw-licking, orofacial grooming and body grooming stage (Fig. 1B, and Movie S1). Then, we explored the role of PF nucleus in mediating self-grooming. By using RNAscope analysis, we found that the PF neurons are predominantly *Cbln2*^+^ (more than 98% across 3 different sections, sFig.1A-B). To measure the contribution of *Cbln2*^+^ PF neurons to self-grooming, we injected AAV-DIO-ChR2-mCherry into the PF of Cbln2-IRES-Cre mice, followed by optical fiber implantation above the PF bilaterally (Fig. 1C). The effect of photostimulation to trigger action potential firing in PF neurons expressing ChR2 was validated in acute brain slices (Fig. 1D). Optogenetic activation (473□nm, 10 s, 2 mW, 20 Hz) of *Cbln2*^+^ PF neurons significantly suppressed spontaneous self-grooming (sFig.1C-J), including paw-licking (Fig. 1E), orofacial grooming (Fig. 1F), and body grooming (Fig. 1G) in mice (Movie S2). Furthermore, optogenetic activation of *Cbln2*^+^ PF neurons also significantly suppressed oil-induced self-grooming (Xie et al., 2022) (Fig. 1H). These data suggested a preliminary conclusion that optogenetic activation of *Cbln2*^+^ PF neurons inhibit both spontaneous and oil-induced self-grooming.

**FIG. 1.**
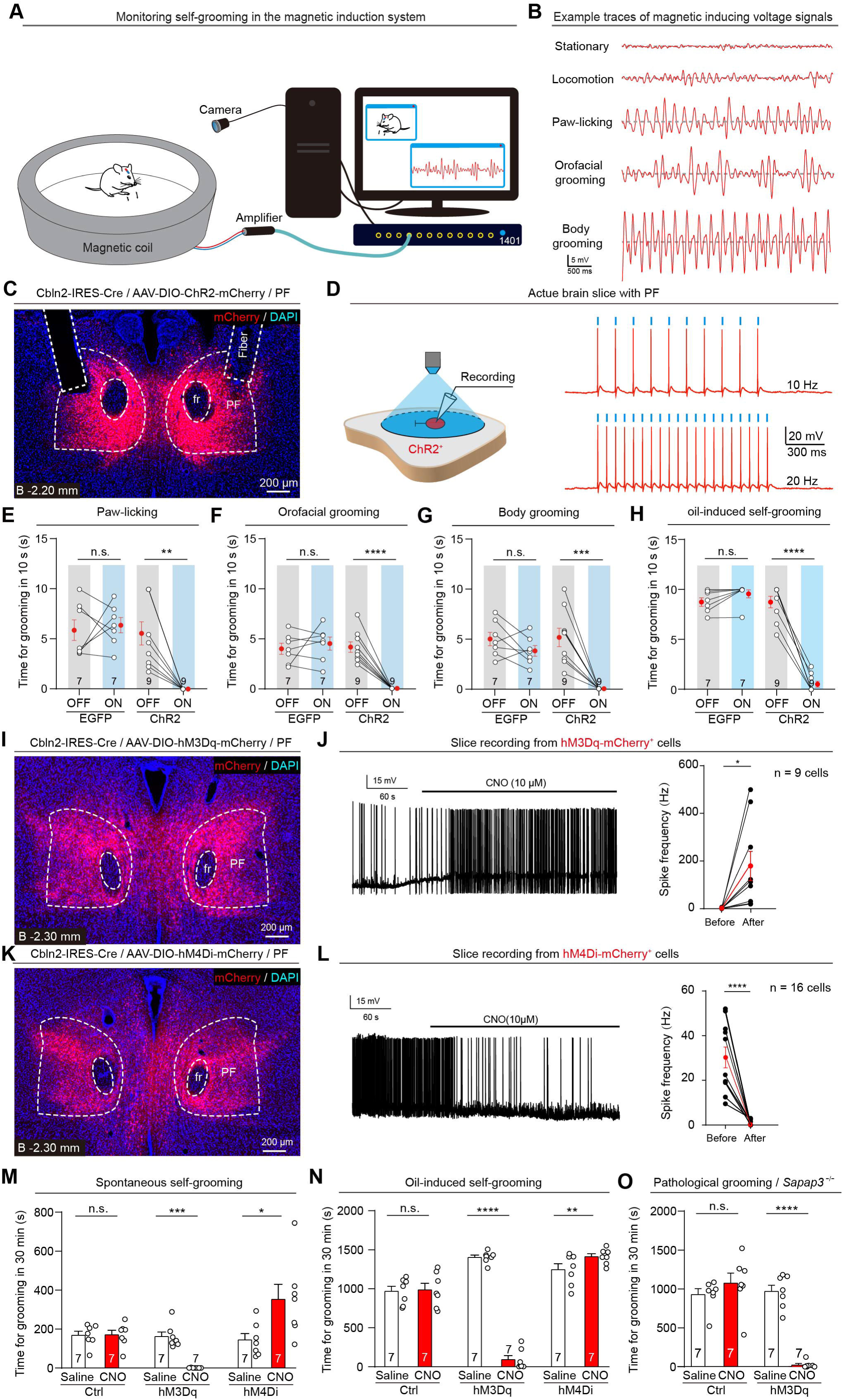
Cbln2-expressing PF neurons are essential for self-grooming. (**A**, **B**) Schematic diagram (**A**) and example voltage pulses traces (**B**) showing the magnetic induction system in parallel with a high-speed camera to record the stationary state, locomotion, paw-licking, orofacial grooming and body grooming in mice. (**C**) Example coronal section showing ChR2-mCherry expression in PF of Cbln2-IRES-Cre mice and optical fiber tracks above the PF. (**D**) Light-pulse trains (473 nm, 10 s, 2 mW, 20 Hz / 10 Hz) reliably evoked phase-locked spiking activity in ChR2-mCherry^+^ PF cells. (**E**-**G**) Quantitative analyses of time spent for self-grooming within 10 second before (OFF) and during (ON) photostimulation of *Cbln2*^+^ PF neurons. Photostimulation was conducted when the mice were in the sequential phases of paw-licking (**E**), orofacial grooming (**F**), and body grooming (**G**). (**H**) Quantitative analyses of time spent for oil-induced self-grooming of mice within 10 second before (OFF) and during (ON) photostimulation of *Cbln2*^+^ PF neurons. (**I**) Example coronal section showing bilateral injection of AAV-DIO-hM3Dq-mCherry into the PF of Cbln2-IRES-Cre mice. (**J**) Example trace of action potential firing (left) and quantitative analysis of firing rate (right) showing the effectiveness of CNO to chemogenetically activate hM3Dq-expressing PF neurons in acute brain slices. CNO was dissolved in artificial cerebrospinal fluid (ACSF) (10 μM) and perfused to the brain slices. (**K**) Example coronal section showing bilateral injection of AAV-DIO-hM4Di-mCherry into the PF of Cbln2-IRES-Cre mice. (**L**) Example trace of action potential firing (left) and quantitative analysis of firing rate (right) showing the effectiveness of CNO to chemogenetically silence hM4Di-expressing PF neurons in acute brain slices. CNO was dissolved in artificial cerebrospinal fluid (ACSF) (10 μM) and perfused to the brain slices. (**M-N**) Quantitative analyses of time spent for spontaneous self-grooming (**M**) and oil-induced self-grooming (**N**) within 30 min in mice treated with saline or CNO (1 mg/kg) to chemogenetically activate or inactivate *Cbln2*^+^ PF neurons. (**O**) Quantitative analyses of time spent for pathological self-grooming within 30 min in mice treated with saline or CNO (1 mg/kg) to chemogenetically activate *Cbln2*^+^ PF neurons in Cbln2-IRES-Cre / *Sapap3*^-/-^ mice. Scale bars and numbers of mice are indicated in the graphs. Data in (**E** - **H**), (**J**), (**L**), (**M**) and (**N**) are means ± SEM (error bars). Statistical analyses in (**E** - **H**), (**J**), (**L**), (**M**) and (**N**) were performed using Student’s t-test (\**p* < 0.05, \*\**p* < 0.01, \*\*\**p* < 0.001, and \*\*\*\**p* < 0.0001). For *p* values, see Supplementary Table 1.

Next, we injected AAV-DIO-hM3Dq-mCherry or AAV-DIO-hM4Di-mCherry into the PF of Cbln2-IRES-Cre mice to verify the function of *Cbln2*^+^ PF neurons in regulation of grooming behavior within a long period (Fig. 1I, K). The effect of CNO on chemogenetical modulation of action potential firing was validated (Fig. 1J, L). We found that chemogenetic activation of *Cbln2*^+^ PF neurons significantly inhibited spontaneous self-grooming, including paw-licking (Fig. 1E), orofacial grooming (Fig. 1F), and body grooming (Fig. 1M and sFig.2B, D), as well as oil-induced self-grooming (Fig. 1N), whereas chemogenetic inhibition of these neurons enhanced both spontaneous and oil-induced self-grooming (Fig. 1M-N and sFig.2C, D), suggesting that *Cbln2*^+^ PF neurons regulate grooming behavior in a bidirectional manner and potentially play key role in the maintenance of balance in the grooming behavior. At the other hand, chemogenetic activation of *Cbln2*^+^ PF neurons caused slight reduction of total distance, while chemogenetic inhibition of *Cbln2*^+^ PF neurons showed no effect on total locomotion distance (sFig.2A). Besides, optogenetic activation of *Cbln2*^+^ PF neurons also did not significantly change the total distance or immobile time (sFig.1K, L), revealing that the regulation of *Cbln2*^+^ PF neurons on grooming behavior was not caused by the obvious motor system dysfunction.

The *Sapap3*-knockout (*Sapap3*^-/-^) mouse is a model used to study OCD, as it recapitulates OCD-like compulsive behavior through excessive grooming (sFig.2E) (Hadjas et al., 2020; Welch et al., 2007). We surprisingly found that chemogenetic activation of *Cbln2*^+^ PF neurons also strongly inhibit pathological self-grooming in *Sapap3*^-/-^ mice (Fig. 1O). These data suggested that *Cbln2*^+^ PF neurons are important components in the central circuit module that broadly controls physiology and pathological self-grooming in mice.

### *Cbln2*^+^ PF neurons encode self-grooming

Next, we systematically characterized the physiological properties of *Cbln2*^+^ PF neurons. To assess the response properties of these neurons during self-grooming, we recorded GCaMP signals in *Cbln2*^+^ PF neurons using fiber photometric recording. AAV-DIO-GCaMP7s was injected into the PF of Cbln2-IRES-Cre mice, followed by implantation of an optical fiber above the *Cbln2*^+^ PF neurons (Fig. 2A). Because the spontaneous grooming behavior does not frequently occur, we sprayed the corn oil over the mice from head to toe to induce more grooming behavior. In most trials, we found the mice exhibited three major types of behavior, including stationary state, locomotion and grooming.

**FIG. 2.**
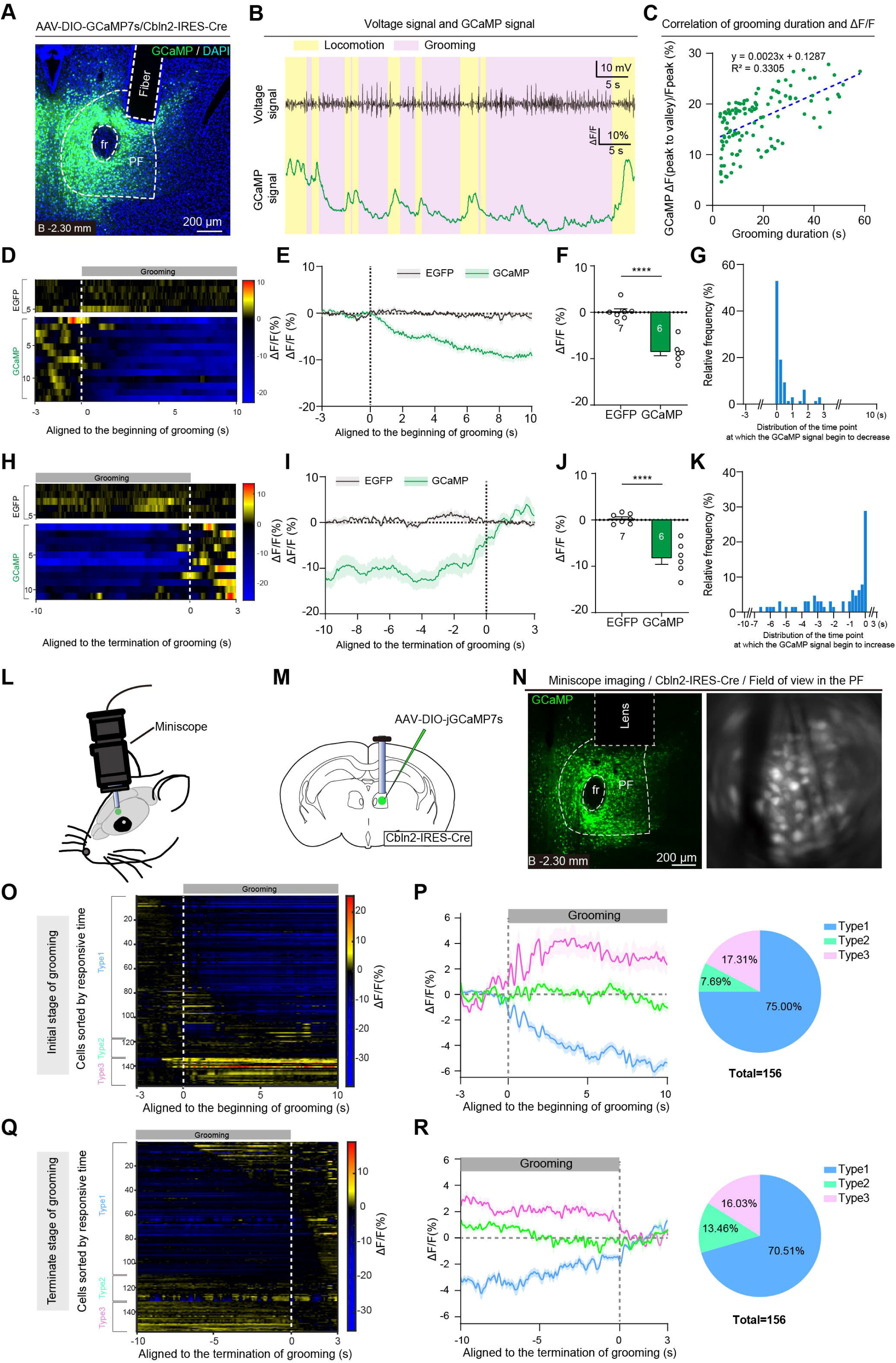
*Cbln2*-expressing PF neurons respond to self-grooming. (**A**) Representative micrographs showing the expression of GCaMP7s in PF, and the optical fiber track above the GCaMP^+^ PF neurons for fiber photometric recording in adult Cbln2-IRES-Cre mice. (**B**) Example GCaMP signals and voltage pulses traces from magnetic induction system in parallel with self-grooming. (**C**) Correlation analysis of average GCaMP7s fluorescence change (ΔF_(peak-valley)_/F_peak_) with grooming duration. (**D**, **H**) Heat-map graphs of GCaMP7s fluorescence of the *Cbln2*^+^ PF neurons aligned to the onset (**D**) and the offset (**H**) of self-grooming. EGFP were used as control. (**E**, **I**) GCaMP response curve aligned with the initiation (**E**) and termination (**I**) of self-grooming of an example mouse, showing kinetics of GCaMP signal of *Cbln2*^+^ PF neurons. EGFP were used as control. (**F**, **J**) Quantitative analyses of average GCaMP7s fluorescence change (ΔF/F) with the initiation (**F**) and termination (**J**) of self-grooming. (**G**, **K**) Quantitative analyses of distribution of the time point at which the GCaMP signal begin to decrease during the initial stage of grooming (**G**) and increase during the terminate stage of grooming (**H**). (**L**) Schematic diagram showing the schedule for recording GCaMP7s fluorescence from *Cbln2*^+^ PF single neurons by miniscope. (**M**) Schematic diagram showing viral injection and GRIN lens implantation strategy for calcium imaging of *Cbln2*^+^ PF neurons. (**N**) Example coronal brain section showing the track of GRIN lens above the GCaMP^+^ PF neurons in Cbln2-IRES-Cre mice (left). The field of view of raw GCaMP signals (right). (**O**) Heatmaps of the activities (ΔF/F0) in individual PF neurons (n = 6 mice). The vertical white dashed line denotes the grooming behavior onset (0 s). The aligned traces were sorted by the responsive time, and classified to 3 different types depending on the average GCaMP fluorescence intensity difference between the grooming period and the baseline period. Type1: decrease; Type2: no change; Type3: increase. (**P**) Average GCaMP fluorescence response (ΔF/F0) curves of 3 different types of neurons aligned with the initiation of grooming behavior (left). Proportion of PF neurons from each category (right, n = 156 neurons from 6 mice). (**Q**) Heatmaps of the activities (ΔF/F0) in individual PF neurons (n =6 mice). The vertical white dashed line denotes the end of grooming behavior (0 s). The aligned traces were sorted by the responsive time, and classified to 3 different types depending on the average GCaMP fluorescence intensity difference between the grooming period and the baseline period. Type1: increase; Type2: no change; Type3: decrease. (**R**) Average fluorescence response (ΔF/F0) curves of 3 different types of neurons aligned with the termination of grooming behavior (left). Proportion of PF neurons from each category (right, n = 156 neurons from 6 mice). Scale bars and numbers of mice are indicated in the graphs. Data in (**E**, **F**) and (**I**, **J**) are means ± SEM (error bars). Statistical analyses in **(F)** and **(J)** were performed using Student’s t-test (\*\*\*\**p* < 0.0001). For *p* values, see Supplementary Table 1.

GCaMP fluorescence of *Cbln2*^+^ PF neurons during grooming was higher than that of during stationary state, and lower than that of during locomotion (sFig. 3A, B). Considering that the grooming behavior is a kind of motor-related behavior, we chose to define the GCaMP fluorescence during the period of locomotion as the baseline. These results lead to a conclusion that *Cbln2*^+^ PF neurons negatively regulate the grooming behavior, which is consistent with the results that opotogenetic or chemogenetic activation of *Cbln2*^+^ PF inhibited grooming behavior.

In parallel with the oscillation of magnetic inducing voltage signal which indicated the period of grooming, the GCaMP fluorescence of *Cbln2*^+^ PF neurons decreased immediately after the grooming behavior occurred, and stayed fairly low until grooming stopped. (Fig. 2B) We also noticed that the relative GCaMP fluorescence of *Cbln2*^+^ PF showed negative correlation with the grooming duration (grooming duration is defined as the duration of a bout, pauses longer than 3 s constitute a new bout), suggesting that the *Cbln2*^+^ PF neurons might also be involved in the maintenance of grooming. (Fig. 2C).

To further analyze the temporal relationship between GCaMP signal and self-grooming, we averaged the trials by aligning the GCaMP signal with the exact time points of initiation or termination of self-grooming. The GCaMP signal decreased immediately after the initiation of self-grooming (Fig. 2D-F). We defined the offset time of GCaMP signal as the time when the signal reaches 5% valley amplitude relative to the baseline. The average time when GCaMP signal began to decrease was 329 ± 71 ms after the initiation of self-grooming (Fig. 2G). The GCaMP signal began to recover to the baseline in prior to the termination of self-grooming (Fig. 2H-J). We defined the onset time of GCaMP signal as the time when the signal recovers to 95% valley amplitude relative to the baseline. The average time when GCaMP signal began to rise was 2531 ± 336 ms before the termination of self-grooming (Fig. 2K). These results suggest that *Cbln2*^+^ PF neurons are more likely involved in the initiation, maintenance and termination of self-grooming.

To explore the neuronal activity of *Cbln2*^+^ PF neurons at the single-cell level, we performed in vivo calcium imaging by using the miniscope system on free-moving mice (see methods for details, Fig. 2L). AAV-DIO-GCaMP7s was injected into PF, followed by the implantation of a gradient refractive index (GRIN) lens above the PF two weeks after viral injection (Fig. 2M). The example coronal brain section revealed the site of GCaMP expression was restricted to the PF, and the track of lens was exactly above the PF (Fig. 2N). In the trials that recorded the initial stage of grooming (Movie S2), we detected three types of GCaMP fluorescence response in *Cbln2*^+^ PF neurons corresponding to the grooming behavior (Fig. 2O, sFig. 3C). 75.00% of *Cbln2*^+^ PF neurons (Type 1) showed decreased response, 17.31% of *Cbln2*^+^ PF neurons (Type 3) showed increased response and 7.61% of *Cbln2*^+^ PF neurons (Type 2) remained unchanged (Fig. 2P), consistent with the results of fiber photometric recording that the total response of GCaMP fluorescence was decreased during grooming compared with that during baseline period (Fig. 2D-F). In the trials that recorded the terminate stage of grooming (Movie S2), the GCaMP fluorescence response in *Cbln2*^+^ PF neurons could also be classified into three types (Fig. 2Q, sFig. 3D). Most of the *Cbln2*^+^ PF neurons (70.51%, Type 1) showed increased response, 16.03% of *Cbln2*^+^ PF neurons (Type 3) showed deceased response and 13.46% of *Cbln2*^+^ PF neurons (Type 2) remained unchanged when the grooming behavior terminated (Fig. 2R), which was also consistent with the results of fiber photometric recording which recorded the terminate stage of grooming (Fig. 2H-J). Together, these data further confirmed the conclusion that *Cbln2*^+^ PF neurons play a negative role in the regulation of self-grooming.

### Mapping the monosynaptic inputs of *Cbln2*^+^ PF neurons

Using recombinant rabies virus, we next performed monosynaptic retrograde tracing (Wickersham et al., 2007) to examine how *Cbln2*^+^ PF neurons are connected to brain areas associated with self-grooming (Fig. 3A–C). A brain-wide survey revealed a series of monosynaptic projections to the *Cbln2*^+^ PF neurons (Fig. 3D, E). Among these brain regions, we focused on the brain regions such as SC, SNr, ZI and LPB. First, robust monosynaptic inputs arose from the superior colliculus (SC) (Fig. 3D2). Prenatal or early postnatal factors could disrupt SC functions, thus resulting in autism (Jure, 2019; Liu et al., 2022), which was reported to show excessive grooming behavior in rodents (Peça et al., 2011). Second, *Cbln2*^+^ PF neurons were monosynaptically innervated by neurons in the zona incerta (ZI) and substantia nigra pars reticulata (SNr) (Fig. 3D3, D4), two brain areas that are involved in the self-grooming behavior in mice (Jiang et al., 2024; Meyer-Luehmann et al., 2002; Zhu et al., 2025). Third, *Cbln2*^+^ PF neurons were also innervated by neurons in the lateral parabrachial nucleus (LPB) (Fig. 3D1). Previous study showed that LPB glutamatergic (Glu) neurons were involved in stress-induced self-grooming (Jia et al., 2023). These data supported the hypothesis that *Cbln2*^+^ PF neurons are within a brain network for self-grooming behavior.

**FIG. 3.**
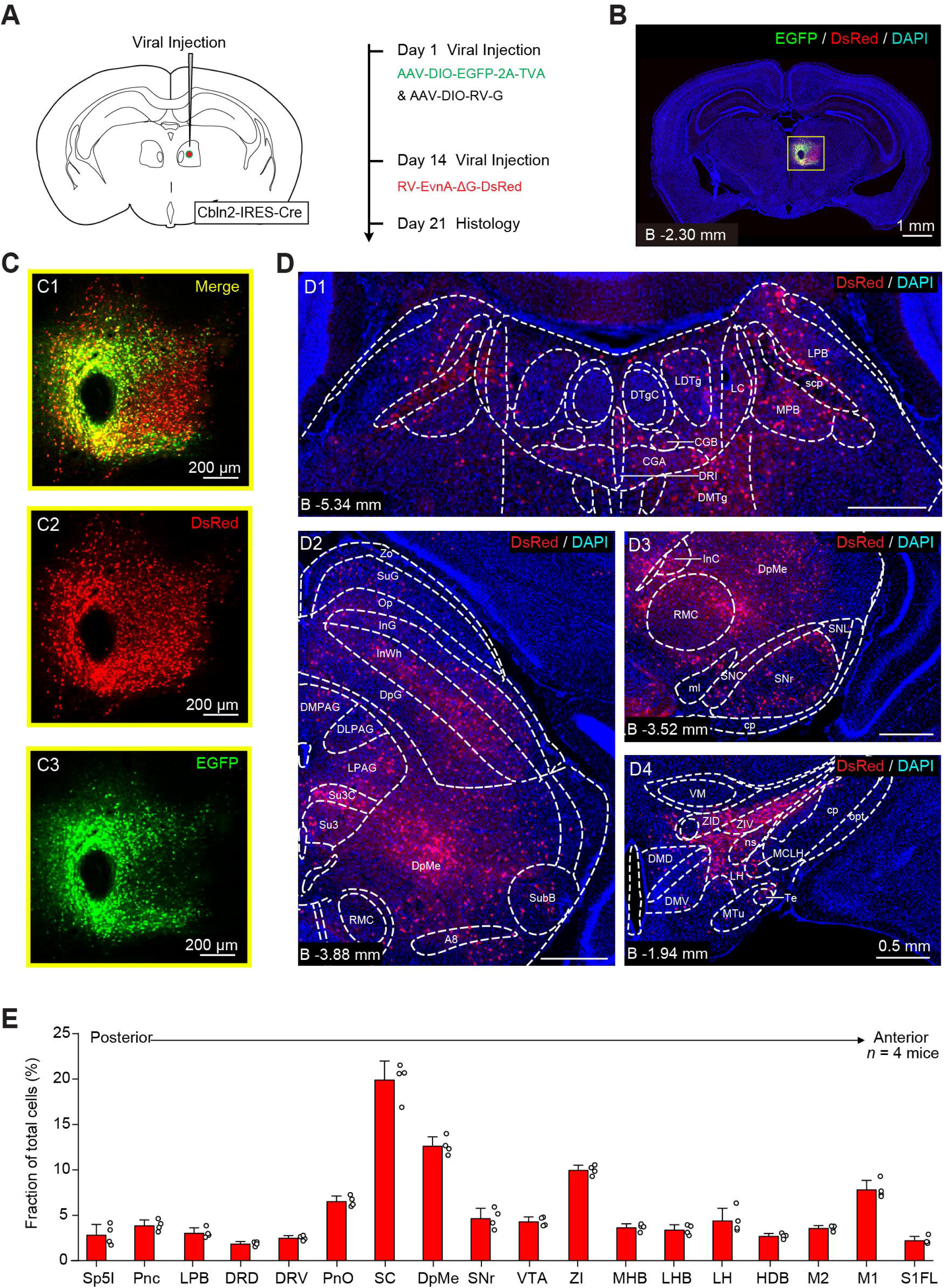
Retrograde tracing of *Cbln2*^+^ PF neurons with RV(A) Schematic diagram showing the viral injection strategy for the retrograde tracing of *Cbln2*^+^ PF neurons using a combination of AAV and rabies virus (RV). (**B-C**) Representative coronal section showing that the viral injection center, as indicated by the co-expression of EGFP and DsRed, was localized in the PF. Note that the double-labeled cells indicate starter cells **(C)**. **(D)** Example micrographs showing DsRed^+^ cells in different brain regions, including the LPB (D1), the SC (D2), the SNr (D3) and the ZI (D4). **(E)** Fraction of total RV-labeled cells in different brain regions monosynaptically projecting to the *Cbln2*^+^ PF neurons. Note that data in **(E)** were normalized by dividing the total number of DsRed+ cells in these brain regions. Numbers of mice **(E)** are indicated in the graphs. Scale bars are labeled in the graphs. Data in **(E)** are means ± SEM (error bars). The experiment **(B-D)** was repeated four times independently with similar results.

### Multiple *Cbln2*^+^ PF neurons input pathways involved in self-grooming

Next, we tested whether projections from multiple brain regions to *Cbln2*^+^ PF neurons were involved in the regulation of self-grooming. We injected AAV-DIO-hM3Dq- mCherry into the SC (Fig. 4A, B) or LPB (Fig. 4E, F) of vGluT2-IRES-Cre mice, which followed by cannulae implantation above the PF (Fig. 4C, G). By the perfusion of CNO into the PF through the cannulae, the vGluT2^+^ SC or LPB-PF pathway was activated. We found that chemogenetic activation of either vGlut2^+^ SC-PF pathway (Fig. 4D) or vGlut2^+^ LPB-PF pathway (Fig. 4H) significantly inhibited spontaneous self-grooming, including paw-licking, orofacial grooming, and body grooming in mice (sFig. 4B, C). Then, we injected AAV-DIO-hM3Dq-mCherry into the SNr (Fig. 4I, J) or ZI (Fig. 4M, N) of vGAT-IRES-Cre mice, which followed by cannulae implantation above the PF (Fig. 4K, O). We found that chemogenetic activation of either vGAT^+^ SC-PF pathway (Fig. 4L) or vGAT^+^ ZI-PF pathway (Fig. 4P) significantly facilitated spontanous self-grooming, including paw-licking, orofacial grooming, and body grooming in mice (sFig. 4D, E). These results verified the inhibitory role of the glutamatergic inputs from SC or LPB to *Cbln2*^+^ PF neurons and the excitatory role of the GABAergic inputs from SNr or ZI to *Cbln2*^+^ PF neurons on grooming behavior, further elucidating that *Cbln2*^+^ PF neurons are within a complex brain network and bidirectionally regulate grooming behavior.

**FIG. 4.**
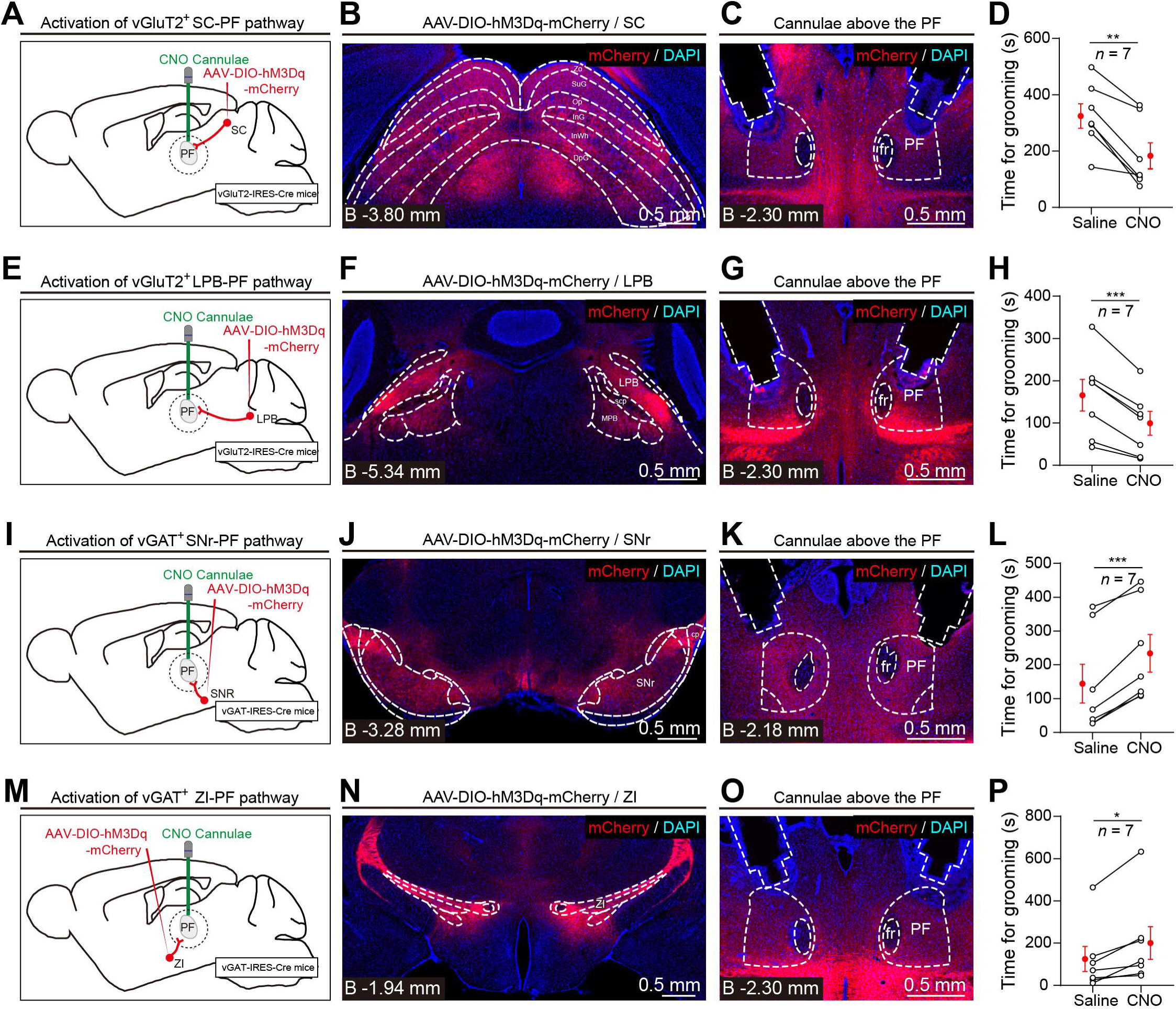
*Cbln2*^+^ PF input pathways involvement in self-grooming. (**A, E**) Schematic diagram showing the injection of AAV-DIO-hM3Dq-mCherry into the SC or LPB of vGluT2-IRES-Cre mice, followed by cannulae implantation above the PF to enable cell-type-specific activation of the vGluT2^+^ SC–PF pathway (**A**) or vGluT2^+^ LPB–PF pathway (**E**). (**B-C, F-G**) Example micrographs showing hM3Dq-mCherry expression in the SC (**B**) or LPB (**F**) and the cannulae above hM3Dq-mCherry^+^ axon terminals in the PF (**C**, **G**). (**D, H**) Quantitative analyses of time spent for spontaneous self-grooming within 30 min in mice treated with saline or CNO (1 mg/kg) to chemogenetically activate vGluT2^+^ SC–PF pathway (**D**) or vGluT2^+^ LPB–PF pathway (**H**). (**I, M**) Schematic diagram showing the injection of AAV-DIO-hM3Dq-mCherry into the SNr or ZI of vGAT-IRES-Cre mice, followed by cannulae implantation above the PF to enable cell-type-specific activation of the vGAT^+^ SNr–PF pathway (**I**) or vGAT^+^ ZI–PF pathway (**M**). (**J-K**, **N-O**) Example micrographs showing hM3Dq-mCherry expression in the SNr (**J**) or ZI (**N**) and the cannulae above hM3Dq-mCherry^+^ axon terminals in the PF (**K**, **O**). (**L, P**) Quantitative analyses of time spent for spontaneous self-grooming within 30 min in mice treated with saline or CNO (1 mg/kg) to chemogenetically activate vGAT^+^ SNr–PF pathway (**L**) or vGAT^+^ ZI–PF pathway (**P**). Scale bars and numbers of mice are indicated in the graphs. Data in (**D**), (**H**), (**L**), and (**P**) are means ± SEM (error bars). Statistical analyses in (**D**), (**H**), (**L**), and (**P**) were performed using Student’s t-test (\**p* < 0.05, \*\**p* < 0.01, and \*\*\**p* < 0.001). For *p* values, see Supplementary Table 1.

### The *Cbln2*^+^ PF–striatum pathway is required for self-grooming

Next, we injected AAV-DIO-EGFP into the PF of Cbln2-IRES-Cre mice to map the efferent pathways of *Cbln2*^+^ PF neurons involved in the grooming behavior. (Fig. 5A-B). We found that the *Cbln2*^+^ PF neurons divergently projected to various brain regions, including the dorsal striatum (CPu), and the ventral striatum (aca, AcbC and Tu). (Fig. 5C), and other brain regions (sFig. 5A-H). We tested the striatum as the possible relay station mediating PF-related grooming behavior as following reasons: first, the activity of striatal cells is reported to be orchestrated to represent the important timepoints in a grooming sequence (Minkowicz et al., 2023); second, striatal dysfunction and defects in cortico-striatal function have been observed in repetitive and compulsive grooming behaviors in autism model mice (Peça et al., 2011) and striatal-specific expression of *Sapap3* rescues the over-grooming behavior in OCD mice (Welch et al., 2007); third, ventral capsule/ventral striatum DBS significantly reduced OCD symptoms in human (Graat et al., 2017; Welter et al., 2021).

**FIG. 5.**
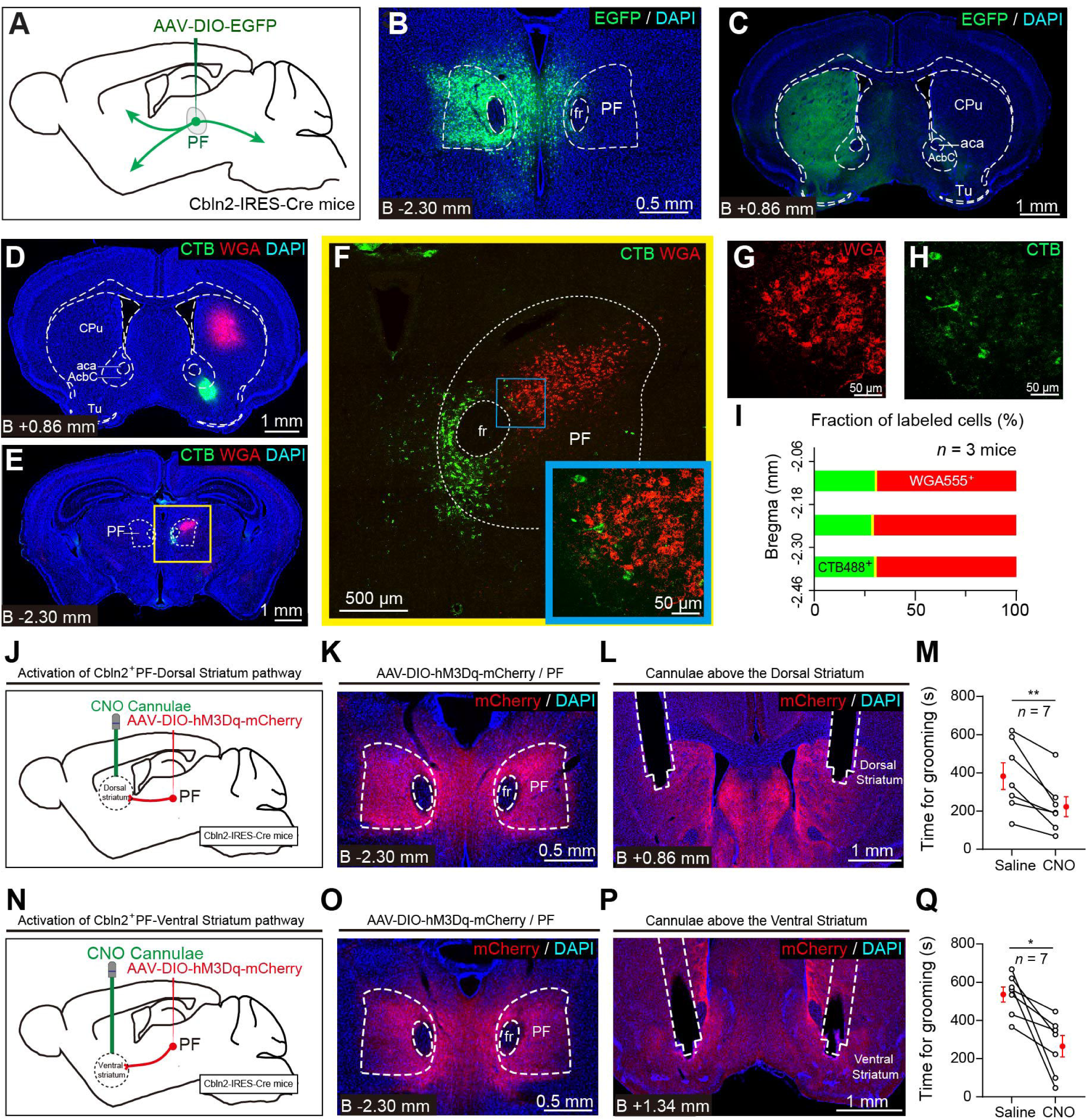
The role of *Cbln2*^+^ PF–striatum pathway in self-grooming. (**A**) Schematic diagram showing the viral injection strategy for labeling *Cbln2*^+^ PF neurons using AAV-DIO-ChR2-mCherry in adult Cbln2*-*IRES-Cre mice. (**B-C**) Representative micrographs from the PF showing EGFP expression in the PF neurons (**B**) and EGFP-positive axon terminals in the striatum (**C**). (**D**) Retrograde labelling of PF neurons by injecting WGA555 into dorsal striatum and CTB488 into ventral striatum. (**E**-**H**) Example coronal brain sections showing dorsal striatum-projecting WGA^+^ PF neurons and ventral striatum-projecting CTB+ neurons. (**I**) Quantitative analyses reflecting the overlap between dorsal striatum- and ventral striatum-projecting PF neurons within different bregmas, showing that dorsal striatum-projecting and ventral striatum-projecting *Cbln2*+ PF neurons are largely segregated. (**J, N**) Schematic diagram showing the injection of AAV-DIO-hM3Dq-mCherry into the PF of Cbln2-IRES-Cre mice, followed by cannulae implantation above the dorsal striatum or ventral striatum to enable cell-type-specific activation of the *Cbln2*^+^ PF–dorsal striatum pathway (**J**) or *Cbln2*^+^ PF–ventral striatum pathway (**N**). (**K-L, O-P**) Example micrographs showing hM3Dq-mCherry expression in the PF (**K**, **O**) and the cannulae above hM3Dq-mCherry^+^ axon terminals in the dorsal striatum (**L**) or ventral striatum (**P**). (**M**, **Q**) Quantitative analyses of time spent for spontaneous self-grooming within 30 min in mice treated with saline or CNO (1 mg/kg) to chemogenetically activate *Cbln2*^+^ PF–dorsal striatum pathway (**M**) or *Cbln2*^+^ PF–ventral striatum (**Q**). Scale bars and numbers of mice are indicated in the graphs. Data in (**M**) and (**Q**) are means ± SEM (error bars). Statistical analyses (**M** and **Q**) were performed using Student’s t-test (\**p* < 0.05, and \*\**p* < 0.01). For *p* values, see Supplementary Table 1.

The striatum, consisting of dorsal and ventral part, receives inputs from distinct subpopulations of neurons in PF (Zhang et al., 2022). The ventral part of the striatum and the ventral striatal islands of Calleja were particularly reported to be involved in the regulation of grooming behavior in rodents (Xue et al., 2022; Zhang et al., 2021), however, function of the dorsal part of striatum in grooming behavior was poorly elucidated. Hence it is essential to verify if PF regulates grooming behavior through two distinct PF-striatum pathways.

By using the strategy of retrograde labeling of cholera toxin subunit B (CTB) and wheat germ agglutinin (WGA) (Fig. 5D), we labelled PF neurons which sent projections to the dorsal striatum and the ventral striatum (Fig. 5E). The dorsal striatum-projecting PF neurons and ventral striatum-projecting PF neurons were scarcely overlapped (double labelled neurons were no more than 1.5%) in spatial arrangement (Fig. 5F-I). It would be essential to clarify the role of these two distinct pathways in PF-mediating grooming behavior regulation. Therefore, we injected AAV-DIO-hM3Dq- mCherry into the PF of Cbln2-IRES-Cre mice (Fig. 5J, K, N, O), which followed by cannulae implantation above the dorsal striatum (Fig. 5L) or ventral striatum (Fig. 5P). We found that chemogenetic activation of either Cbln2^+^ PF-dorsal striatum pathway (Fig. 5M) or Cbln2^+^ PF-ventral striatum pathway (Fig. 4H) significantly inhibited spontaneous self-grooming, including paw-licking, orofacial grooming, and body grooming in mice (sFig. 4F, G). Together, these data elucidated the pivotal role of both the dorsal striatum and ventral striatum in the regulation of PF-mediating grooming regulation.

In addition to the majority of GABAergic spiny projection neurons, there are many kinds of interneurons in the striatum, which act on nearby circuits and shape functional output to the basal ganglia (Muñoz-Manchado et al., 2018). Optogenetic activation of parvalbumin-positive striatal interneurons reduced self-grooming events in *Sapap3*^-/-^ mice (Mondragón-González et al., 2024). By the injection of AAV-DIO-EGFP-Syb2 into the PF of Cbln2-IRES-Cre mice (sFig.5I), we found that the striatal cells positive for Parvalbumin (PV), Somatostatin (SST), Choline Acetyltransferase (ChAT) and Neuropeptide Y (NPY) in both the dorsal striatum and ventral striatum were surrounded by the synaptic terminals from *Cbln2*^+^ PF neurons (sFig. 5J, K), which offered putative targets to divergently regulate the grooming behavior. However, further work was essential to elucidate the functional connections between *Cbln2*^+^ PF neurons and these striatal cells and the role of these striatal cells in grooming regulation under the innervation of *Cbln2*^+^ PF neurons.

### Ultrasound stimulation targeting the PF results in activation of PF neurons and alleviates over-grooming in OCD mice

Activation of *Cbln2^+^* PF neurons significantly reduces time spent for grooming in both WT mice and OCD mice (Fig. 1E-H, M-O), and nearly all the neurons are Cbln2 positive in PF (sFig. 1A, B), making PF an ideal target for therapeutic intervention. Therefore, we explored whether non-invasive approach, such as ultrasound stimulation, would efficiently activate PF neurons and have impact on time spent for spontaneous grooming in WT or OCD mice.

We conducted a preliminary experiment to determine whether ultrasound stimulation targeting the PF could affect spontaneous grooming time in WT mice (Fig. 6A, B). Under the schedule of 3 days of habituation followed by 7 days of daily 30-minute ultrasound stimulation, we monitored and compared the grooming duration of mice after habituation (Day 3) or each daily stimulation session (Day 4 – Day 10, sFig. 6A). We found a slight decrease in spontaneous grooming time on Day 6, with significant reductions observed on Day 8, 9, and 10. Therefore, in the formal experiment, we chose to apply daily 30-minutes stimulation for 7 days and assessed grooming behavior only on the Day 3 (for baseline) and Day 10 (Fig. 6C). We found that ultrasound stimulation targeting the PF not only increased c-Fos expression in the PF region (Fig 6D, E), but also enhanced neuronal excitability of PF neurons (Fig 6F, G). As expected, it also reduced spontaneous grooming duration in WT mice (Fig 6H, I). Using the same stimulation schedule, we applied ultrasound stimulation to OCD mice and observed a significant reduction in their excessive grooming behavior (Fig 6J, K). Although the grooming time was not reduced to WT levels, this non-invasive approach still offers promising potential as a therapeutic strategy for OCD.

**FIG. 6.**
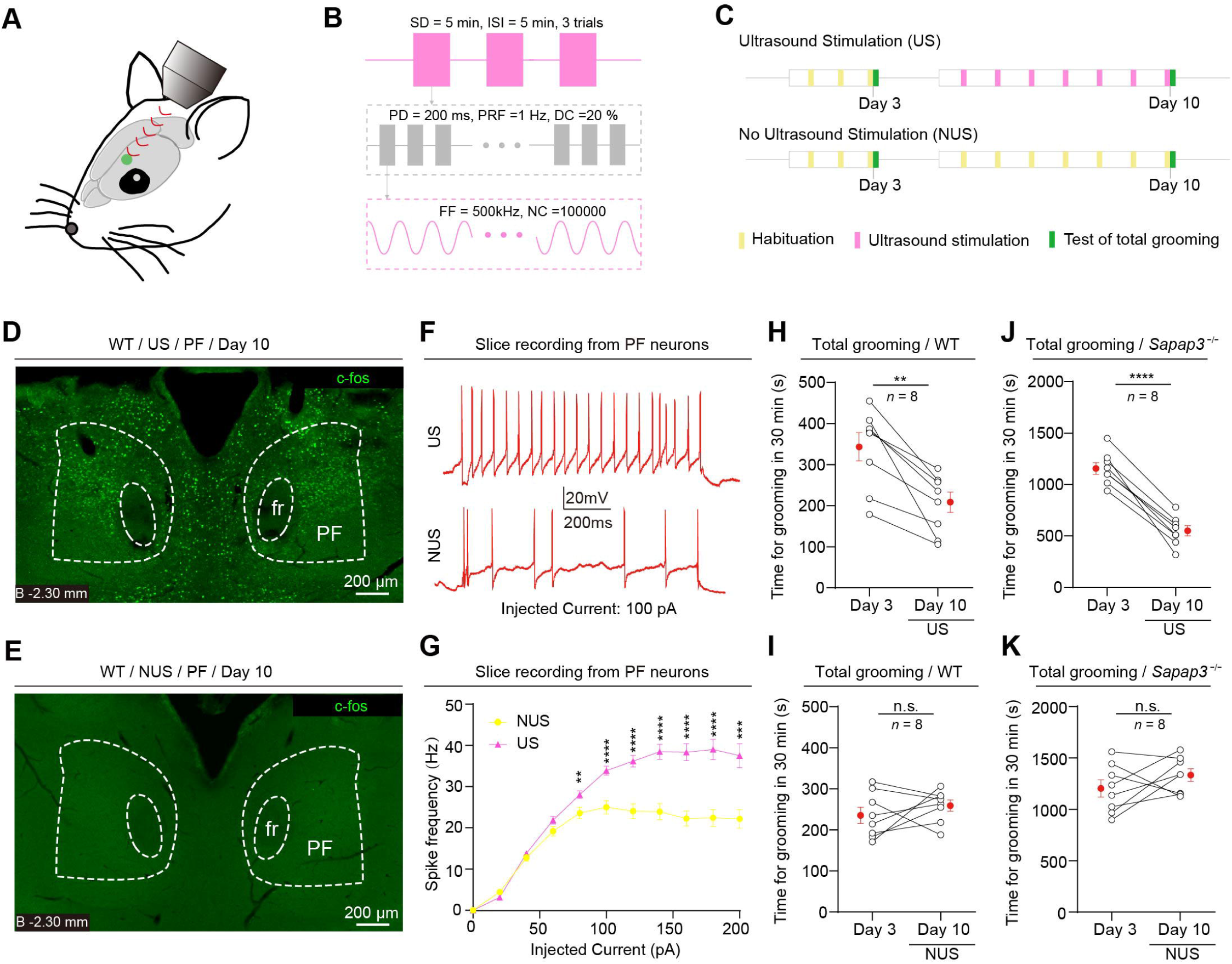
Ultrasound stimulation targeting PF alleviates over-grooming in *Sapap3^-/-^* mice. (**A**) Schematic of ultrasound stimulation. (**B**) Parameters of ultrasonic stimulation, inter-stimulus interval (ISI), fundamental frequency (FF), pulse repetition frequency (PRF), number of cycle(NC), duty cycle (DC), pulse duration (PD) and stimulation duration (SD) of the ultrasound signal. (**C**) Schematic diagram showing the schedule of ultrasound stimulation and recording of grooming for test mice and control mice. (**D**, **E**) Example coronal section showing c-fos staining of PF in the test mouse with ultrasound stimulation (US) (**D**) and control mouse with no ultrasound stimulation (NUS) (**E**). (**F**) Example trace of depolarization-induced spiking activity in PF neurons from test mice and control mice, Injected current = 100 pA. (**G**) Quantitative analysis of the firing rates in PF neurons from test mice and control mice in response to step depolarizing current injections. (**H**, **I**) Quantitative analyses of time spent for spontaneous grooming of WT mice (**H**) and *Sapap3*^-/-^ mice (**I**) on Day 3 (baseline) and Day 10 (with 30-min ultrasound stimulation from day 4 to day 10 for each day). (**J**, **K**) Quantitative analyses of time spent for spontaneous grooming of WT mice (**J**) and *Sapap3*^-/-^ mice (**K**) on Day 3 (baseline) and Day 10 (only habituation for 30 mins from day 4 to day 10 for each day). Scale bars and numbers of mice are indicated in the graphs. Data in (**G** - **K**) are means ± SEM (error bars). Statistical analyses in (**H** - **K**) were performed using Student’s t-test (\**p* < 0.05, \*\**p* < 0.01, \*\*\**p* < 0.001, and \*\*\*\**p* < 0.0001). For *p* values, see Supplementary Table 1.

## DISCUSSION

Self-grooming is an innate behavior with important physiological functions including hygiene maintenance, thermoregulation, social communication (Kalueff et al., 2016; Leonard et al., 2005; Spruijt et al., 1992). Recently, the repetitive stereotyped behavior of mice was reported to cope with stress and post-stress anxiety alleviation (Sun et al., 2022). In this study, we identified that *Cbln2*^+^ PF neurons encode self-grooming in a bidirectional manner. Synaptic excitatory inputs that activate *Cbln2*^+^ PF neurons inhibit spontaneous self-grooming, and synaptic inhibitory inputs enhance it. Moreover, *Cbln2*^+^ PF neurons regulate self-grooming behavior through both dorsal and ventral striatum. Together, these data elucidate that *Cbln2*^+^ PF neurons are a central component of a brain network involved in self-grooming behavior.

Our data also showed that chemogenetic activation of *Cbln2*^+^ PF neurons substantially decreased the excessive pathological self-grooming in OCD mice (Welch et al., 2007), providing a solid neurobiological foundation for the improvement of OCD/OCB symptoms under the treatment of DBS in PF in human (Servello et al., 2020). Meanwhile, our preliminary findings indicated that activating PF neurons through non-invasive ultrasound stimulation could also reduce excessive grooming behavior in OCD mice, which offers additional options for clinical treatment of OCD.

The PF nucleus participates in sensorimotor coordination (Liu et al., 2008), pain and nociception (Sims-Williams et al., 2016; Weigel and Krauss, 2004). Sensory perception is one of the cases inducing grooming behavior (Xie et al., 2022). Among the brain regions we mapped which send inputs to PF, SC and LPB are reported to mediate the circuits of sensory perception (Choi et al., 2023; Nagashima et al., 2023), and ZI is believed to have many integrative functions that link sensory stimuli with motor responses to guide behavior (Hormigo et al., 2023). Optogenetic and chemogenetic activation of *Cbln2*^+^ PF neurons inhibit the oil-induced self-grooming, reveal the importance of the PF nucleus in sensory-induced grooming behavior, while may also suggest that the PF nucleus is broadly involved in the regulation of grooming behavior within a complex neural network.

*Cbln2*^+^ PF neurons send projections to various brain regions, including the dorsal striatum, the ventral striatum and others. Here we have proven that PF-dorsal striatum pathway and PF-ventral striatum pathway are spatially independent but both involved in self-grooming regulation, specifying the effect of dorsal striatum on grooming behavior which was rarely reported in rodents. Besides the striatum receiving outputs from *Cbln2*^+^ PF neurons, the periaqueductal gray (PAG) and the ventral tegmental area (VTA) are also potentially innervated by *Cbln2*^+^ PF neurons and reported to implicate in grooming behavior (Gao et al., 2019; Sun et al., 2022). In future studies, it would be important to examine whether these two brain regions are involved in PF-mediating grooming regulation, thereby enriches the landscape of research on the regulation of grooming behavior.

There are certain limitations in our present study. First, the magnetic inducing voltage signals induced by embedding a magnet in the head only provide a relatively more comprehensive reflection of grooming behavior. However, further analysis on the characteristics of the voltage signals corresponding to different types of grooming will be helpful to automatically distinguish among them. Second, *Cbln2* is expressed in the vast majority of PF neurons, with the proportion exceeding 98%. Our miniscope data revealed three types of neurons with different responses to grooming behavior, addressing the question that classification of detailed subtypes of neurons in PF nucleus combining with single-cell sequence would be necessary to better elucidate how PF neurons encode self-grooming. Third, chemogenetic activation of *Cbln2*^+^ PF neurons significantly alleviated the excessive grooming in OCD mice, while further electrophysiological, structural and biochemical studies are essential to refine the neural mechanism of *Cbln2*^+^ PF neurons on pathological grooming regulation, which would prospectively bring new therapeutic target for OCD. Finally, we have only preliminary confirmed that ultrasound stimulation targeting PF reduces grooming time, future studies are required to determine how long the activation of PF neurons induced by ultrasound stimulation persist and the optimal stimulation parameters to prolong or enhance the activation effects.

## ACKNOWLEDGMENTS

This work was supported by STI 2030-Major Projects [Grant 2021ZD 0203200-04 (to F.Z.)], National Natural Science Foundation of China [Grant 32171018 (to F.Z.), Grant 32271030 (to D.L.)], Natural Science Foundation of Hebei Province [Grant H2022206011 (to F.Z.), C2024206032 (to F.Z.)], Top Talent of Young People of Hebei Province (to F.Z.).

## AUTHOR CONTRIBUTIONS

Conceptualization, P.C., F.Z., and L.Z.; methodology, D.L.; investigation, D.G., Y.L., Z.L., H.S., L.W., Z. X., B.Y., L.Z., R.Z., T.Z., and Z.Z.; writing– original draft, P.C., F.Z., and L.Z.; funding acquisition, D.L. and F.Z.; resources, P.C., D.L. and F.Z.; supervision, P.C., F.Z., and L.Z.

## METHODS

### Animals

All the mice-using experimental procedures were conducted following the protocols approved by the Administrative Panel on Laboratory Animal Care in the National Institute of Biological Sciences (NIBS) in Beijing, China. Mouse lines used in this study, including vGluT2-Cre and VGAT-Cre were imported from the Jackson Laboratory (JAX Mice and Services). Cbln2-IRES-Cre mice were generated in NIBS and *Sapap3*^-/-^ mice were purchased from the GemPharmatech Co., Ltd. Mice were housed in groups (3–5 animals per cage) and maintained under a 12-h light/12-h dark cycle with food and water available *ad libitum*. Mice were separated 3 days before virus injection and housed individually after virus injection until subsequent experiments were conducted. Primers used for genotyping are listed in Supplementary Table 2.

### AAV vectors

Viral particles used in this study were purchased from Shanghai Taitool Bioscience Inc., BrainVTA Inc. and OBiO Technology Inc. Detailed information about the AAVs were listed in the Supplementary Table 2. The final titer used for AAV injection was 5×10^12^ viral particles/ml.

### Stereotaxic injection

Mice were anesthetized with an intraperitoneal injection of tribromoethanol (125–250 mg per kg). Standard surgery was performed to expose the brain surface above the target brain regions, including PF, SC, LPB, ZI, SNr, dorsal striatum and ventral straitum. Coordinates used to inject PF were as follows: bregma -2.30 mm, lateral ±0.65 mm and dura -3.20 mm. Coordinates used to inject SC were as follows: bregma -3.80 mm, lateral ±1.20 mm and dura -1.55 mm. Coordinates used for LPB injection were as follows: bregma -5.43 mm, lateral ±1.35 mm and bregma -3.50 mm. Coordinates used for ZI injection were as follows: bregma -1.79 mm, lateral ±1.28 mm and dura -4.09 mm. Coordinates used for SNr injection were as follows: bregma -3.13 mm, lateral ±1.37 mm and dura -4.36 mm. Coordinates used for dorsal striatum injection were as follows: bregma +0.86 mm, lateral ±1.70 mm and dura -2.60 mm. Coordinates used for ventral striatum injection were as follows: bregma +1.20 mm, lateral ±1.50 mm and dura -4.45 mm. The AAVs, CTB-488 and WGA-555 were stereotaxically injected using a glass pipette connected to a Nanoliter-Injector 201 (World Precision Instruments) at a slow flow rate of 0.15 μl min^−1^ to avoid potential damage to local brain tissue. The pipette was withdrawn at least 20 min after viral injection.

For optogenetic activation, AAV injections were bilateral. For chemogenetic activation/inactivation, injections were bilateral. For chemogenetically synaptic activation, injections were bilateral. Behavioral tests were conducted 3 weeks after viral injection. Slice physiology and histology were conducted at least 3 weeks after AAV injection.

### Optical fiber implantation

Thirty minutes after AAV injection, a ceramic ferrule with an optical fiber (for optogenetics: 200 µm in diameter, numerical aperture (NA) of 0.37; for fiber photometric recording: 200 µm in diameter, NA of 0.37) was implanted with the fiber tip on top of the PF (bregma -2.30 mm, lateral ± 1.71 mm and bregma -2.55 mm with a 20° angle towards the midline). The ferrule was then secured onto the skull with dental cement. After implantation, the antibiotics were applied to the surgical wound. The optogenetic and fiber photometric recording experiments were conducted 3 weeks after the optical fiber implantation. For optogenetic stimulation, the output of the laser was measured and adjusted to various intensities of 2 mW or 0.2 mW before each experiment. The pulse onset, duration, and frequency of light stimulation were controlled by a programmable pulse generator attached to the laser system.

### Cannula implantation and drug infusion

Two cannulae were bilaterally implanted above the PF, dorsal striatum or ventral striatum regions. The inner and outer diameters of the cannula were 150 μm and 300 μm, respectively. Each cannula was fixed to the skull with dental cement. During drug infusion, each cannula was connected with a catheter filled with CNO (3 μM) or saline for injection. The other side of the catheter was connected to a Hamilton syringe controlled by an infusion pump to drive the delivery of CNO or saline (50 nl min−1). The catheter was removed 5 minutes after the delivery of CNO or saline was completed. Then the mice were back to homecage for a 30-minute rest and used for the subsequent behavior tests. At the end of the experimental session, the test mice were perfused, and the coronal brain sections containing PF, dorsal striatum or ventral striatum regions were inspected for the presence of cannula track. The mice with no cannula track above the PF, dorsal striatum or ventral striatum were rejected from further analysis.

### Preparation of behavioral tests

After AAV injection and fiber implantation, the mice were housed individually for 3 weeks, and handled daily by the experimenters for at least 3 days before the behavioral tests. On the day of the behavioral test, the mice were transferred to the testing room and were habituated to the room conditions for 3 h before the experiments started. The apparatus was cleaned with 20% ethanol, to eliminate odor cues from other mice. All behavioral tests were conducted during the same circadian period (13:00–19:00). All behaviors were scored by the experimenters, who were blinded to the animal treatments.

### Magnetic induction to record head movements during self-grooming

The magnetic induction system to measure self-grooming was constructed in the light of a magnetic induction system to measure scratching behavior (Inagaki et al., 2002). For the mice without fibers or cannulae, a small skin cut (1 mm) was made in the scalp under anesthesia. A metal guide tube with sharp tip that carried a magnet cylinder (1 mm in diameter, 3 mm in length) was inserted subcutaneously. Then the cylinder was pulled out, while leaving the magnet inside the scalp. A drop of vet glue was applied to help the skin cut sealed well and healed. The mice could be tested 3 days after the surgery. For the mice with fibers or cannulae, the magnet cylinder was placed on the skull and wrapped in dental cement during the surgery of fiber/cannula implantation. Then the behavior tests were conducted 3 weeks after the surgery.

The tested mouse was placed in the chamber in a Helmholtz coil (230 mm in diameter, CTXQ-230, Li Tian magnetoelectric Technology Inc.), or in the arena above a Helmholtz coil. The electric potential induced in the coil by the movement of magnets was amplified (100x) and recorded with Spike2 software (CED Inc., UK). In Spike2 software, the raw electrical potential data was removed DC offset, passed through a digital low pass filter (Butterworth, fourth order, corner: 40 Hz) and smoothed (time constant: 0.04 s). The mean voltage of when the mice were stationary was calculated as the baseline. The threshold to identify the head movement during self-grooming was defined as the mean voltage of baseline plus 5 x SD (standard deviation).

### Recording the time spent for spontaneous self-grooming and oil-induced self-grooming

On the day of experiment, the test mouse was allowed to explore a cylinder (15 cm in diameter) for 15 minutes. For oil-induced grooming, the experimenter sprayed corn oil on the test mice from the top of the cylinder to make the mice covered with a thin layer of oil after the freely exploration in the cylinder. The behavior of self-grooming was identified by both visual inspection and the magnetic induction trace reflecting the rhythmic head movements. The oil-induced self-grooming was quantified by measuring the time spent for self-grooming during the 30 minutes after the spray of oil onto the whole body of the test mice.

### Fiber photometric recording

A fiber photometric system (ThinkerTech) was used to record GCaMP signals from *Cbln2*^+^ PF neurons. To induce fluorescence signals, a laser beams from a laser tube (488 nm) was reflected by a dichroic mirror, focused by a 10 × (NA of 0.37) lens and coupled to an optical commutator. A 2-m optical fiber (200 μm in diameter, NA of 0.37) guided the light between the commutator and the implanted optical fiber. To minimize photobleaching, the power intensity at the fiber tip was adjusted to 0.02 mW. The GCaMP7s fluorescence was band-pass filtered (MF525-39, Thorlabs) and collected by a photomultiplier tube (R3896, Hamamatsu). An amplifier (C7319, Hamamatsu) was used to convert the photomultiplier tube current output to voltage signals, which were further filtered through a low-pass filter (40 Hz cut-off; Brownlee 440). The analog voltage signals were digitized at 100 Hz and recorded by a Power 1401 digitizer and Spike2 (CED).

AAV-hSyn-DIO-GCaMP7s was stereotacically injected into the PF of Cbln2-IRES-Cre mice followed by the implantation of an optical fiber above the *Cbln2*^+^ PF neurons and the implantation of magnet cylinder inside the dental cement. Three weeks after AAV injection, fiber photometric recording was used to record GCaMP signals from *Cbln2*^+^ PF neurons of free moving mice in the magnetic induction system. A flasing LED triggered by 1s square-wave pluse was simultaneously recorded to synchronize the video, GCaMP signals and the magnetic inducing voltage signals. After the experiments, the optical fiber tip sites in the PF were histologically examined in each mouse.

### Cell-type-specific RV tracing

A modified RV-based three-virus system was used for mapping whole-brain inputs to *Cbln2*^+^ PF neurons (Wickersham et al., 2007). All viruses, including AAV2/9-CAG-DIO**-**EGFP-2A-TVA (5 × 10^12^ viral particles per ml), AAV2/9-CAG-DIO-RVG (5 × 10^12^ viral particles/ml), and EnvA-pseudotyped, glycoprotein (RG)-deleted, and EGFP-expressing RVs (RV-EvnA-ΔG-DsRed) (5.0 × 10^8^ viral particles/ml), were packaged and provided by BrainVTA Inc. (Wuhan, China). A mixture of AAV2/9-CAG-DIO-EGFP-2A-TVA and AAV2/9-CAG-DIO-RVG (1:1, 300 nl) was injected into the PF of Cbln2-IRES-Cre mice unilaterally. Two weeks after AAV helper injection, RV-EvnA-ΔG-DsRed (300 nl) was injected into the same location in the PF of Cbln2-IRES-Cre mice in a biosafety level-2 lab facility. Starter neurons were characterized by the co-expression of DsRed and EGFP, which were restricted in the PF. One week after injection of RV, the mice were perfused with saline, followed by 4% paraformaldehyde (PFA) in phosphate-buffered saline (PBS). The brain was post-fixed in 4% PFA for 8 h and then incubated in PBS containing 30% sucrose until they sank to the bottom. Coronal brain sections (40-μm thickness) were prepared using a cryostat (Leica CM1900). All sections were collected and stained with 4’,6-diamidino-2-phenylindole (DAPI). The sections were imaged with an Olympus VS120 epifluorescence microscope (10× objective lens) and analyzed with ImageJ software. For quantification of subregions, boundaries were based on the Allen Institute’s reference atlas. We selectively analyzed retrogradely labeled dense areas. The fractional distribution of total cells labeled with RV was measured.

### Cell-type-specific anterograde tracing

For cell-type-specific anterograde tracing of *Cbln2*^+^ PF neurons, AAV-DIO-EGFP-syb2 (300 nl) was stereotaxically injected into the PF of Cbln2-IRES-Cre mice. The mice were then maintained in a cage individually. Three weeks after viral injection, mice were perfused with saline followed by 4% PFA in PBS. The brains were post-fixed in 4% PFA for 8 hours and then incubated in PBS containing 30% sucrose until they sank to the bottom. Coronal brain sections at 40 μm in thickness were prepared using a cryostat (Leica CM1900). All coronal sections were collected and stained with primary antibody against GFP and DAPI. The coronal brain sections were imaged with an Olympus VS120 epifluorescence microscope (10 × objective lens).

### Slice physiological recording

Slice physiological recording was performed according to a previously published protocol. Brain slices containing the PF were prepared from adult mice anesthetized with isoflurane before decapitation. Brains were rapidly removed and placed in ice-cold oxygenated (95% O_2_ and 5% CO_2_) cutting solution (in mM: 228 sucrose, 11 glucose, 26 NaHCO_3_, 1 NaH_2_PO_4_, 2.5 KCl, 7 MgSO_4_ and 0.5 CaCl_2_). Coronal brain slices (400 μm) were cut using a vibratome (VT 1200S, Leica Microsystems). The slices were incubated at 28°C in oxygenated artificial cerebrospinal fluid (ACSF containing in mM: 119 NaCl, 2.5 KCl, 1 NaH_2_PO_4_, 1.3 MgSO_4_, 26 NaHCO_3_, 10 glucose and 2.5 CaCl_2_) for 30 min, and then kept at room temperature under the same conditions for 1 h before transfer to the recording chamber. The ACSF was perfused at 1ml min^−1^. The acute brain slices were visualized using a ×40 Olympus water immersion lens, differential interference contrast (DIC) optics (Olympus) and a CCD camera. Patch pipettes were pulled from borosilicate glass capillary tubes (Warner Instruments, no. 64-0793) using a PC-10 pipette puller (Narishige). For recording of action potentials (current clamp), pipettes were filled with a solution (in mM: 135 K-methanesulfonate, 10 HEPES buffer, 1 EGTA (ethylene glycol tetraacetic acid), 1 Na-GTP, 4 Mg-ATP and 2% neurobiotin (pH7.4)). The resistance of pipettes varied between 3.0 and 3.5 MΩ. The voltage signals were recorded using MultiClamp 700B and Clampex 10 data acquisition software (Molecular Devices). After establishment of the whole-cell configuration and equilibration of the intracellular pipette solution with the cytoplasm, series resistance was compensated to 10–15 MΩ. An optical fiber (200μm in diameter) was used to deliver light pulses, with the fiber tip positioned 500 μm above the brain slices. Light-evoked action potentials from ChR2-mCherry^+^ neurons in the PF were triggered by a light-pulse train (473 nm, 2 ms, 10 Hz or 20 Hz, 5 mW) synchronized with Clampex 10 data acquisition software (Molecular Devices).

### Miniscope calcium imaging and data analysis

#### 1) AAV injection and gradient index (GRIN) lens implantation

For miniscope calcium imaging studies, AAV-hsyn-DIO-jGCaMP7s was microinjected into PF of Cbln2-IRES-Cre mice with following coordinates: anterior-posterior, -2.30 mm; medial-lateral, -0.65 mm; dorsal-ventral, -3.2 mm from dura, then the GRIN lens (0.5 mm in diameter × 6.1 mm in length, Inscopix) was implanted 2 weeks after viral injection. For the implantation, the GRIN lens was connected to the imaging system of miniscope to monitor the GCaMP fluorescence during slowly lowering (∼100 μm/min) to the target brain area. After the lens was placed at the appropriate coordinates to gain the optimal intensity of GCaMP fluorescence and anchored to the skull with dental cement, the silicon adhesive (Kwik-Sil, WPI) was used to cap the lens to avoid damage.

#### 2) Measuring GCaMP fluorescence

Miniscope calcium imaging was conducted by using a miniaturized fluorescence microscopy unit (miniscope) linked to a data acquisition device (Inscopix, nVista). At least one day before behavior test and calcium imaging, awake mice with implanted lens were head-fixed on a circular treadmill and a baseplate was implanted above the GRIN lens to get a suitable field for imaging after mounted by the miniscope. Then the miniscope was replaced by a fake one and the mice were returned to their homecages, which enabled the mice to be habituated to the weight of miniscope.

On the day of behavior test and calcium imaging, two side-view cameras were used to record the grooming behavior. Synchronization between imaging data and grooming behavior was implemented by the video recording the flash of LED light and a Power 1401 digitizer which was used to mark the signal generated by the data acquisition device when each session of imaging data was documented. Imaging data were acquired at a 20 Hz frame rate.

#### 3) Data acquisition and analysis

Imaging data were preliminarily processed by the Inscopix data processing software (version 1.6.0). Then the image data were exported as Tiff files. These Tiff files were further analyzed by CNMF-E code in Matlab for the generation of example videos, or manually analyzed by ImageJ software to read the dynamic changes of GCaMP fluorescence frame by frame for each neuron.

For the trials which recorded the initial stage of grooming, the frame at which the mice started to groom was denoted as 0 s, and normalized GCaMP fluorescence (ΔF/F) was calculated over a baseline period of -3 s to 0 s.

For the trials which recorded the terminate stage of grooming, the frame at which the mice stop grooming was denoted as 0 s, and normalized GCaMP fluorescence (ΔF/F) was calculated over a baseline period of 0 s to 3 s.

### Ultrasound stimulation

On the day prior to experimental procedures, the hair on the top of the mouse’s head was carefully removed using hair removal cream. The mice were anesthetized with 3% isofurane in an induction chamber. After sedation, the mice were placed in a supine position on a warming pad (38 °C) to maintain normothermia, and fixed to the three-dimensional brain stereotaxic instrument by a head bar for ultrasound stimulation keeping in light narcosis with 1% isofurane delivered via 100% O_2_ mask inhalation (Li et al., 2019). The schematic of ultrasound stimulation is shown in Fig. 6A. The ultrasound system used was similar to those used in previous studies (Wang et al., 2024; Zhao et al., 2023). In the experiment, a pulse signal generator (AFG3022C, Tektronix, USA) was used to continuously transmit modulated sine wave electrical signals, and the signals were amplified by a power amplifier (E&I 240 L, ENI Inc., USA) and then further amplified by a focused ultrasound probe (V301-SU, focus [diameter, 25.4 mm; radius of curvature, 40 mm], Olympus, USA) during ultrasound stimulation. A conical collimator (3D-printing, resin material, bottom aperture: 4 mm) filled with ultrasonic coupling gel was fixed between the ultrasonic probe and the mouse head, therefore the ultrasonic waves could penetrate the mouse brain, and the ultrasound pressure was 0.36 MPa. The corresponding spatial-peak and pulse-average intensity values were 4.3 W/cm^−2^ and the spatial-peak and time-average intensity value was 215.8 mW/cm^−2^ (Wang et al., 2024; Zhao et al., 2023). As shown in Fig. 6B, the fundamental frequency (FF), pulse repetition frequency (PRF), number of cycle(NC), duty cycle (DC), stimulation duration (SD), pulse duration (PD) and inter-stimulus interval (ISI) of the ultrasound waves were 500 kHz, 1 Hz, 100000, 20%, 300 s, 200 ms and 300 s, respectively. The stimulation was performed at the same time period every day, and each stimulation was followed by a 5-min pause, for a total of three trials of 30 min (Fig. 6B).

Ultrasound stimulation for test mice (US) was continuously performed for 7 days (Fig. 6C). All procedures for the control mice (NUS) were the same as that of the test mice (US) except that the pulse signal generator of the control mice was turned off. Behavioral testing was performed as described in Fig. 6C and sFig.6A. Briefly, for the preliminary experiment, mice were acclimated to the ultrasound system for 30 minutes daily over three days. Grooming time was tested on the third day (Day 3), then the mice received daily stimulation for 7 consecutive days, with each session lasting 30 min. Grooming time was assessed after each simulation session (Day 4 - Day 10). For the formal experiment, mice were acclimated to the ultrasound system for 30 minutes daily over three days and were recorded for grooming time on Day 3 as baseline. Then the mice received 7 days of daily 30-minute ultrasound stimulation, and were recorded for grooming time on Day 10. Mice were sacrificed on Day 10 immediately after recording. Only mice exhibiting significantly increased c-fos staining in the PF region were included in the group of US for further analysis.

### Immunohistochemistry

Mice were anesthetized with isoflurane and sequentially perfused with saline and PBS containing 4% PFA. Post-fixation of the brain was avoided to optimize immunohistochemistry. Brains were incubated in PBS containing 30% sucrose until they sank to the bottom. Cryostat sections (40 μm) were collected, incubated overnight with blocking solution (PBS containing 10% goat or donkey serum and 0.7% Triton X-100), and then treated with primary antibodies diluted with blocking solution for 3 - 4 h at room temperature. Primary antibodies used for immunohistochemistry are listed in the Supplementary Table 2. Primary antibodies were washed three times with washing buffer (PBS containing 0.7% Triton X-100) before incubation with secondary antibodies (tagged with Alexa Fluor® 488 and Alexa Fluor® 546 dilution 1:500; Invitrogen) for 1 h at room temperature. Sections were then washed three times with washing buffer, stained with DAPI, and washed with PBS, transferred onto Super Frost slides, and mounted under glass coverslips with mounting media.

Sections were imaged with an Olympus VS120 epifluorescence microscope (10 × objective lens) or a Zeiss LSM 800 confocal microscope (20 × and 63 × oil-immersion objective lens). Samples were excited by 405, 488 and 561 nm lasers in sequential acquisition mode to avoid signal leakage. Saturation was avoided by monitoring pixel intensity with Hi-Lo mode. Confocal images were analyzed with ImageJ software.

### RNAscope

Mice were perfused with PBS treated with 0.1% DEPC (Sigma, D5758), followed by DEPC-treated PBS containing 4% PFA (PBS-PFA). Brains were post-fixed in DEPC-treated PBS-PFA solution overnight and then placed in DEPC-treated 30% sucrose solution at 4°C for 30h. Brain sections to a thickness of 20 μm were prepared using a cryostat (Leica, CM3050S) and subsequently hybridized with the Cbln2 probe (Advanced Cell Diagnostics, 428551) following the user manual of the RNAscope kit (Advanced Cell Diagnostics, Doc No. 323100-USM). To visualize the NeuN signals, brain sections were incubated with a primary antibody against NeuN at 4°C for 24 h and then with an Alexa Fluor® 594-conjugated goat anti-rabbit secondary antibody (1:500, A11034, Invitrogen) at room temperature for 2 h. Brain sections were imaged using a Zeiss LSM780 confocal microscope or the Olympus VS120 Slide Scanning System. Detailed information about probe and reagents are listed in the Supplementary Table 2.

### Data quantification and statistical analyses

Data collection and analyses were performed blinded to the conditions of the experiments. For statistical analyses of the experimental data, two-sided Student’s t-test was used. The “n” used for these analyses represents the number of mice or cells.

---

**sFIG. 1.**
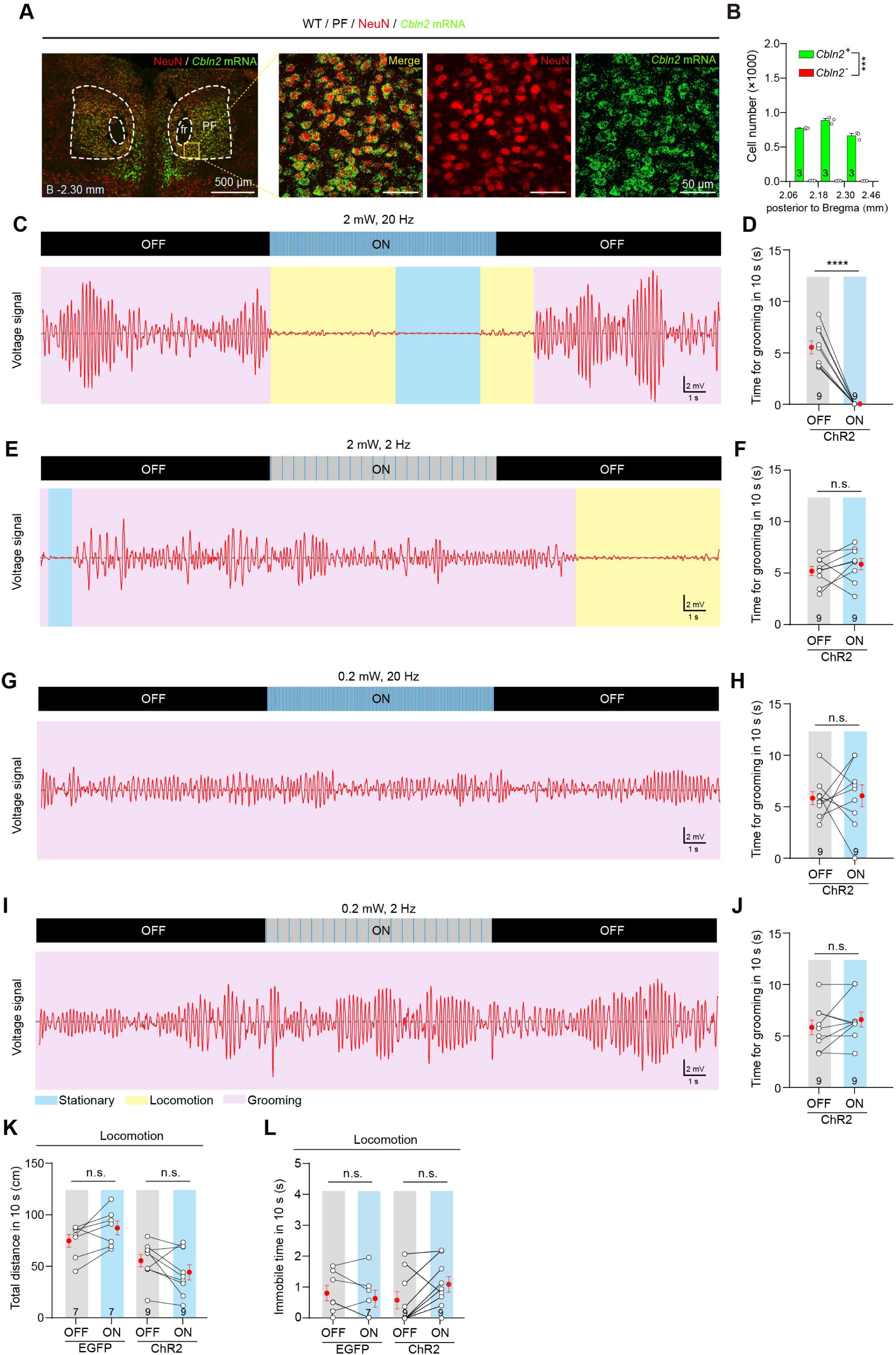
Photostimulation of *Cbln2*^+^ PF neurons inhibit spontaneous self-grooming but locomotion. (**A**) Example micrographs showing the distributions of cbln2 mRNA in the PF of WT mice. (**B**) Quantitative analyses of number of PF cells expressing cbln2 mRNA (green) in coronal sections, as indicated by the distance to bregma. (**C, E, G, I**) Example voltage pulses traces showing the magnetic induction system in parallel with a high-speed camera to record spontaneous self-grooming, within 10 second before (OFF) and during (ON) photostimulation of *Cbln2*^+^ PF neurons by various light-pulse trains (473□nm, 2□ms, 2□mW, 20□Hz) for (**C**), (473□nm, 2□ms, 2□mW, 2□Hz) for (**D**), (473□nm, 2□ms, 0.2□mW, 20□Hz) for (**G**), and (473□nm, 2□ms, 0.2□mW, 2□Hz) for (**I**). (**D, F, H, J**) Quantitative analyses of of time spent for spontaneous self-grooming within 10 second before (OFF) and during (ON) photostimulation of *Cbln2*^+^ PF neurons by various light-pulse trains (473□nm, 2□ms, 2□mW, 20□Hz) for (D), (473□nm, 2□ms, 2□mW, 2□Hz) for (F), (473□nm, 2□ms, 0.2□mW, 20□Hz) for (H), and (473□nm, 2□ms, 0.2□mW, 2□Hz) for (**J**). (**K**, **L**) Quantitative analyses of time for total distance (L) and immobile time (L) of mice within 10 second before (OFF) and during (ON) photostimulation of *Cbln2*^+^ PF neurons. Scale bars are labeled in the graphs. Numbers of mice are indicated in the graphs (**B**, **D**, **F**, **H**, **J**, **K** and **L**). Data in (**B**), (**D**), (**F**), (**H**), (**J**), (**K**) and (**L**) are means ± SEM (error bars). Statistical analyses in (**B**), (**D**), (**F**), (**H**), (**J**), (**K**) and (**L**) were performed using Student’s t-test (\*\*\**p* < 0.001, and \*\*\*\**p* < 0.0001).

**sFIG. 2.**
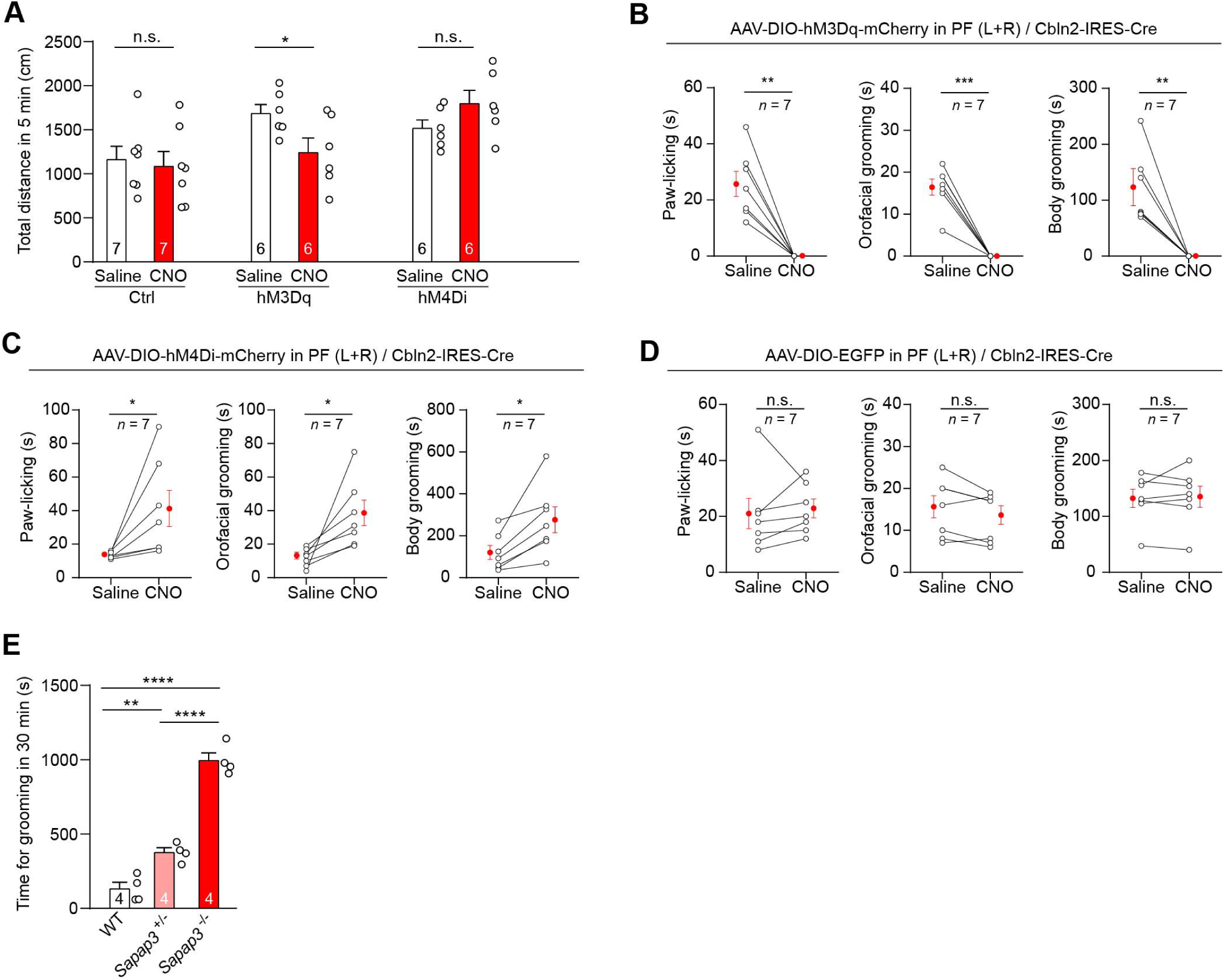
Chemogenetically activation of *Cbln2*^+^ PF neurons inhibit paw-licking, orofacial grooming and body grooming. **(A)** Quantitative analyses of total distance within 5 min in mice treated with saline or CNO (1 mg/kg) to chemogenetically activate or inactivate *Cbln2*^+^ PF neurons. (**B**) Injection of AAV-DIO-hM3Dq-mCherry into PF of Cbln2-IRES-Cre mice, quantitative analyses of time spent for paw-licking, orofacial grooming, and body grooming of mice within 30 min treated with saline or CNO (1 mg/kg) to chemogenetically activate *Cbln2*^+^ PF neurons. (**C**) Injection of AAV-DIO-hM4Di-mCherry into PF of Cbln2-IRES-Cre mice, quantitative analyses of time spent for paw-licking, orofacial grooming, and body grooming of mice within 30 min treated with saline or CNO (1 mg/kg) to chemogenetically inactivate *Cbln2*^+^ PF neurons. (**D**) Injection of AAV-DIO-EYFP into PF of Cbln2-IRES-Cre mice, quantitative analyses of time spent for paw-licking, orofacial grooming, and body grooming of mice within 30 min treated with saline or CNO (1 mg/kg). (**E**) Quantitative analyses of time spent for excessive self-grooming within 30 min in *Sapap3*^+/-^ and *Sapap3*^-/-^ mice. Numbers of mice are indicated in the graphs (**A**-**E**). Data in (**A**-**E**) are means ± SEM (error bars). Statistical analyses in (**A**-**E**) were performed using Student’s t-test (\**p* < 0.05, \*\**p* < 0.01, \*\*\**p* < 0.001, and \*\*\*\**p* < 0.0001).

**sFIG. 3.**
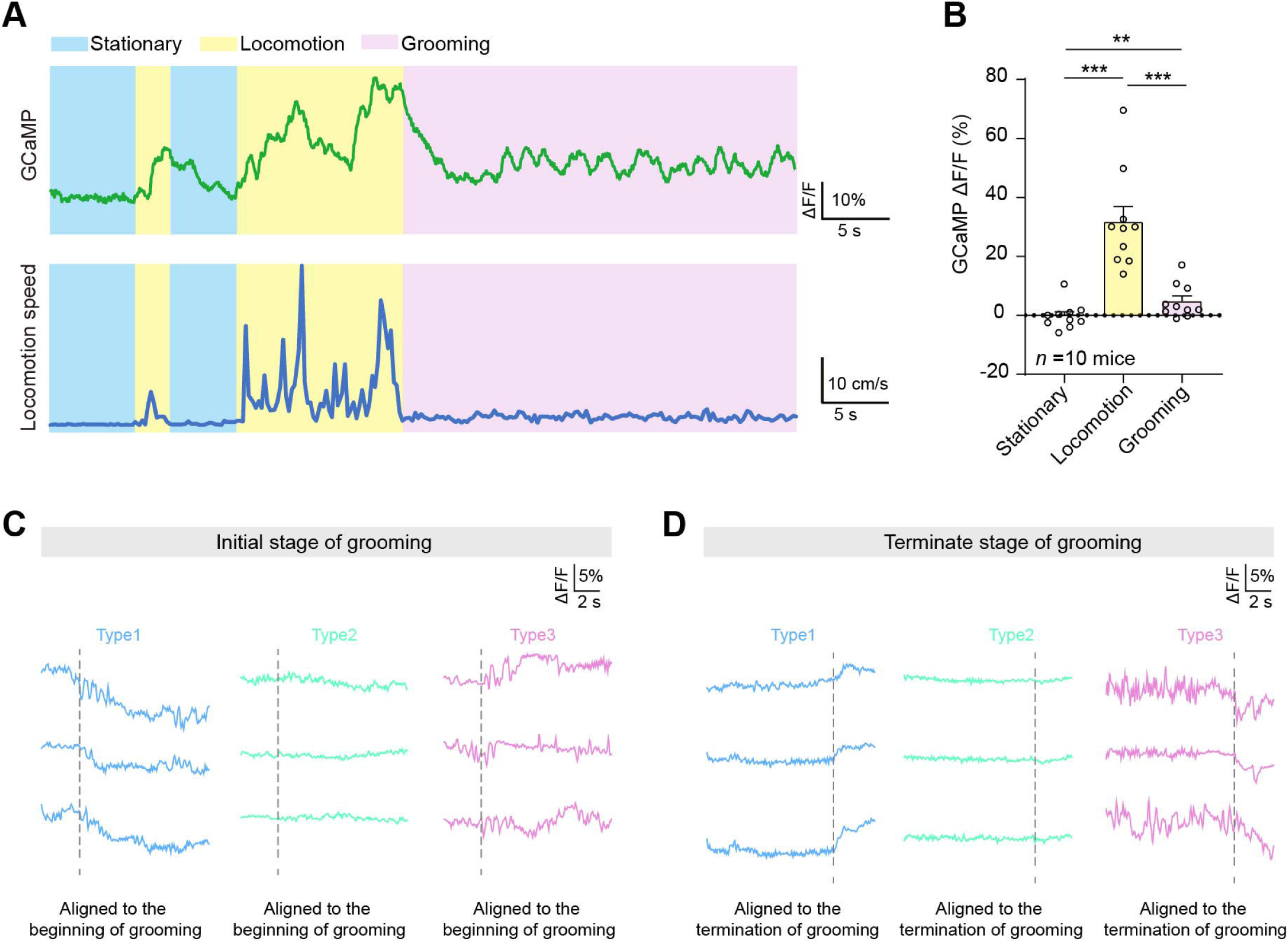
*Cbln2*-expressing PF neurons show decreased Ca^+^ activity during self-grooming. **(A)** Example GCaMP signals from *Cbln2*^+^ PF neurons during the stationary, locomotion, and self-grooming stages by fiber photometric recording. (**B**) Quantitative analyses of average GCaMP7s fluorescence change (ΔF/F) with during the stationary, locomotion, and self-grooming stages. (**C**) Representative GCaMP response curve from single *Cbln2*^+^ PF neurons aligned with the initiation of self-grooming. (**D**) Representative GCaMP response curve from single *Cbln2*^+^ PF neurons aligned with the termination of self-grooming. Numbers of mice are indicated in the graphs (**B**). Data in (**B**) are means ± SEM (error bars). Statistical analyses in (**B**) were performed using Student’s t-test (\*\**p* < 0.01, and \*\*\**p* < 0.001).

**sFIG. 4.**
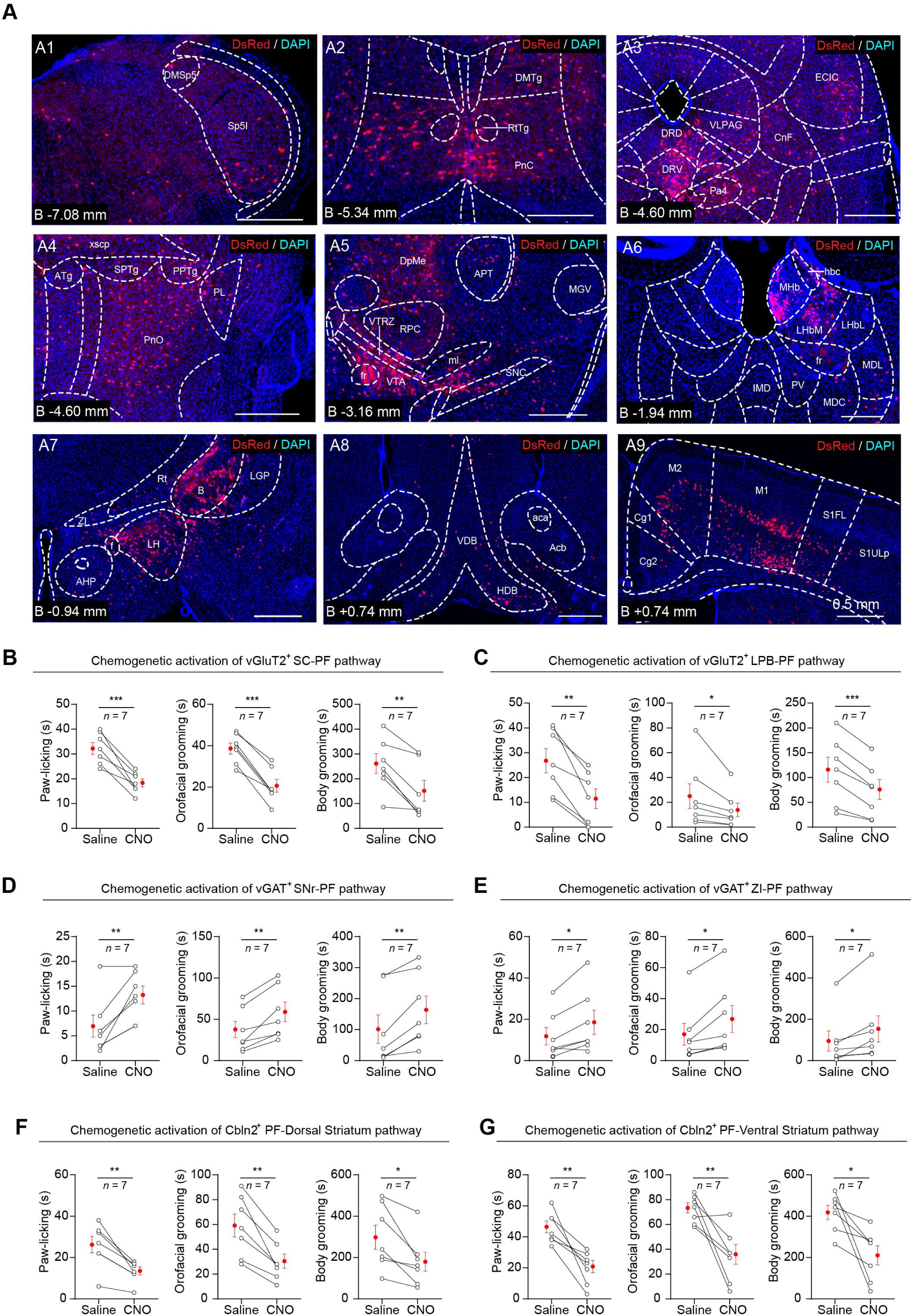
Chemogenetically activation of input pathway of *Cbln2*^+^ PF neurons inhibit paw-licking, orofacial grooming and body grooming. **(A)** Example micrographs showing DsRed^+^ cells in different brain regions, including the Sp5I (A1), the Pnc (A2), the DRD (A3), the DRV (A3), the PnO (A4), the VTA (A5), the MHb (A6), the LH (A7), the HDB (A8), the M1 (A9) and the M2 (A9). Scale bars are labeled in the graphs. Number of mice: four. (**B**-**G**) Quantitative analyses of time spent for paw-licking, orofacial grooming, and body grooming of mice within 30 min treated with saline or CNO (1 mg/kg) to chemogenetically activate vGluT2^+^ SC–PF pathway (**B**), vGluT2^+^ LPB–PF pathway (**C**), vGAT^+^ SNr–PF pathway (**D**), vGAT^+^ ZI–PF pathway (**E**), Cbln2^+^ PF–dorsal striatum pathway (**F**) or Cbln2^+^ PF–ventral striatum (**G**). Numbers of mice are indicated in the graphs (**B**-**G**). Data in (**B**-**G**) are means ± SEM (error bars). Statistical analyses in (**B**-**G**) were performed using Student’s t-test (\**p* < 0.05, \*\**p* < 0.01, and \*\*\**p* < 0.001).

**sFIG. 5.**
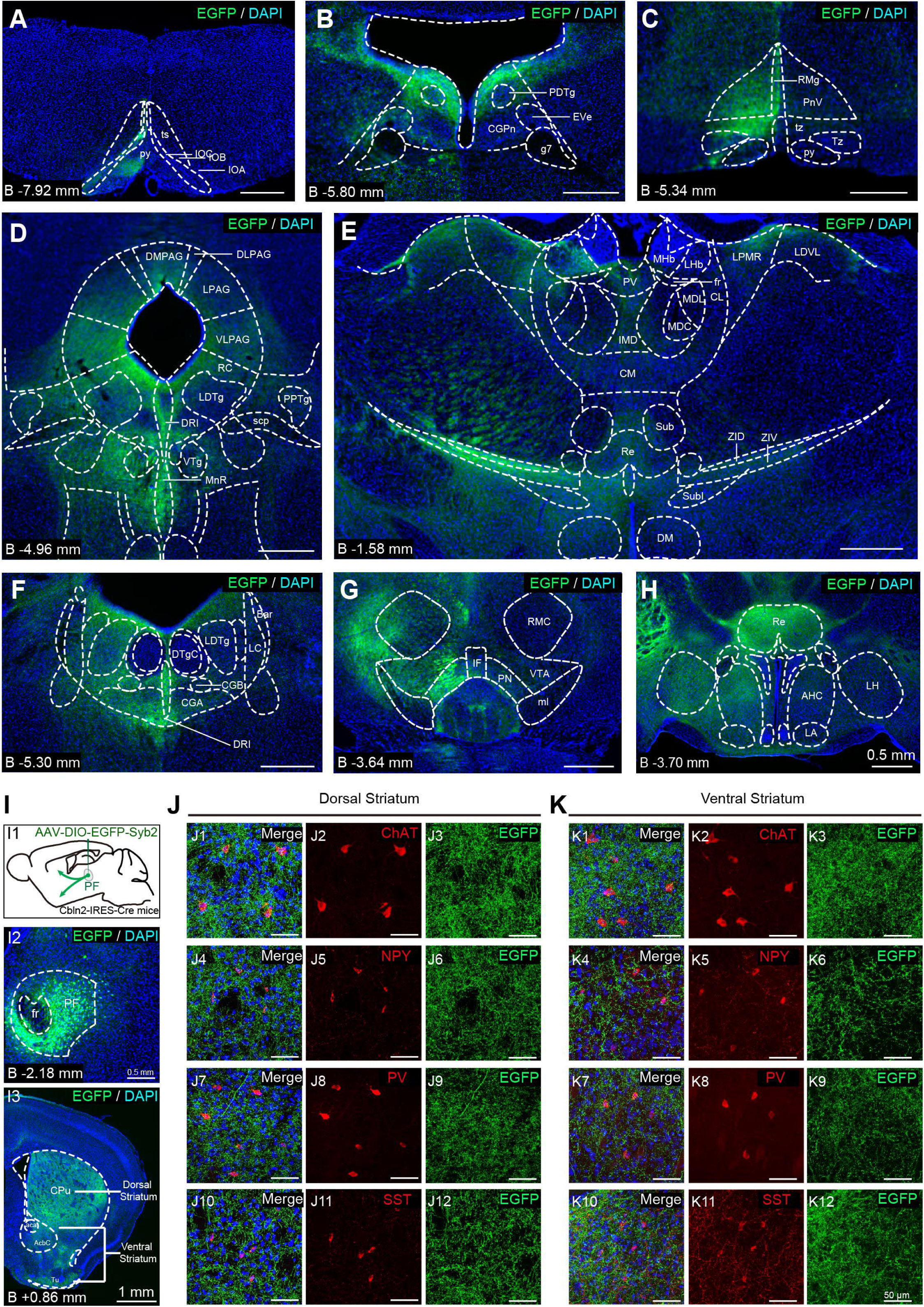
Mapping the monosynaptic outputs of *Cbln2*^+^ PF neurons. (**A**-**H**) Example coronal brain sections showing EGFP^+^ synaptic terminals of *Cbln2*^+^ PF neurons in the target brain regions, including the py (**A**), CGPn (**B**), PnV (**C**), LPAG (**D**), VLPAG (**D**), DRI (**D**), MnR (**D**), ZI (**E**), LC (**F**), CGA (**F**), LDTg (**F**), VTA (**G**), Re (**H**), AHC (**H**), and LH (**H**). (**I**) Antrograde labeling of synaptic terminals of *Cbln2*^+^ PF neurons in the dorsal and ventral striatum. (**J** and **K**) Various subtypes of interneurons, including ChAT^+^, NPY^+^, PV^+^ and SST^+^ in the dorsal stratum (**J**) and ventral stratum (**K**) were surrounded by the EGFP^+^ synaptic terminals from *Cbln2*^+^ PF neurons. Scale bars are labeled in the graphs.

**sFIG. 6.**
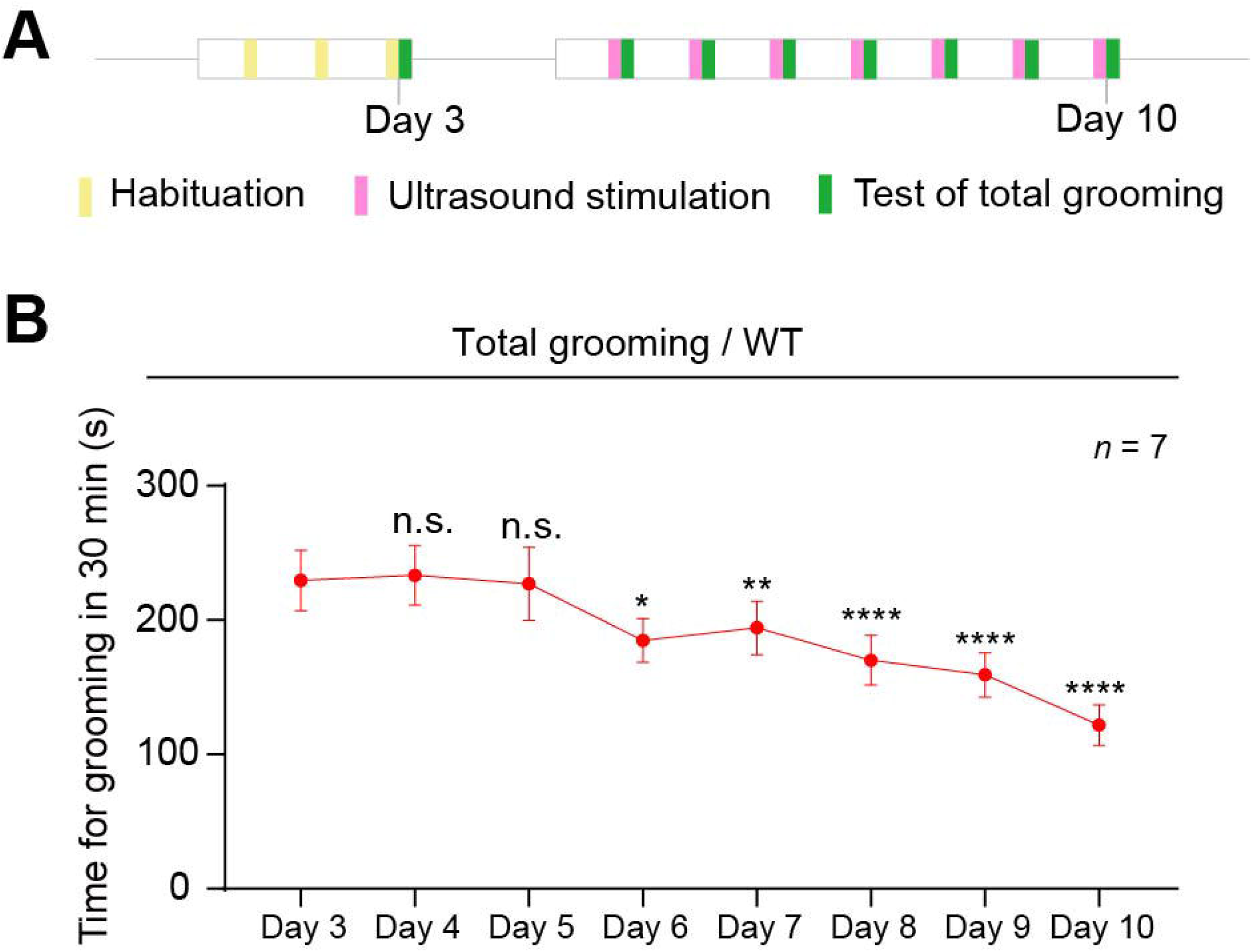
Ultrasound stimulation targeting PF reduces time spent for total grooming in WT mice. (**A**) Schematic diagram showing the schedule for investigation of the effect of ultrasound stimulation on time spent for spontaneous grooming of WT mice. (**B**) Quantitative analyses of time spent for spontaneous grooming of WT mice on each day. Data in (**B**) are means ± SEM (error bars). Statistical analyses in (**B**) were performed using Student’s t-test (\**p* < 0.05, \*\**p* < 0.01, and \*\*\*\**p* < 0.0001).

